# Evolution of resistance and disease tolerance mechanisms to oral bacterial infection in *D. melanogaster*

**DOI:** 10.1101/2023.08.23.554397

**Authors:** Tânia F. Paulo, Priscilla A. Akyaw, Tiago Paixão, Élio Sucena

**Affiliations:** Instituto Gulbenkian de Ciência, Rua da quinta grande 6, 2780-156 Oeiras, Portugal; Departamento de Biologia Animal, Faculdade de Ciências, Universidade de Lisboa, Edifício C2, Campo Grande, 1749-016 Lisbon, Portugal; cE3c: Centre for Ecology, Evolution and Environmental Changes, Faculdade de Ciências, Universidade de Lisboa, Campo Grande, 1749-016 Lisbon, Portugal; CHANGE – Global Change and Sustainability Institute

**Keywords:** Experimental Evolution, Innate Immunity, *Drosophila*, *Pseudomonas entomophila*

## Abstract

Pathogens exert strong selection on hosts, that evolve and deploy different defensive strategies, namely minimizing pathogen exposure (avoidance), directly promoting pathogen elimination (resistance), and/or managing the deleterious effects of illness (disease tolerance). However, how the response to pathogens partitions across these processes has never been directly assessed in a single system, let alone in the context of known adaptive trajectories under controlled selection regimes. Here, an experimental evolution system composed of *D. melanogaster* and its natural pathogen *P. entomophila* is used to independently assess the role of behavioural traits, and of resistance and disease tolerance mechanisms on host evolution. We compare one replicate of a population adapted to oral infection with *P. entomophila* (BactOral) to a replicate of its control population to find no evidence for behavioural change but measurable differences in both resistance and disease tolerance. In BactOral, we identify a relative decrease in bacterial loads correlated with an increase in gut production of specific AMPs, but no differences in bacterial intake, in gut cell renewal rate, or in the rate of bacterial defecation, pointing to a strengthening in resistance. Additionally, we posit that disease tolerance also contributes to the adaptive response of the BactOral population through a tighter control of its immune response and of the deleterious effects of exposure. This study reveals a genetically complex and mechanistically multi-layered response, possibly reflecting the structure of adaptation to infection in natural populations.

## BACKGROUND

Host-parasite interactions are main drivers of evolution^1,2^. In natural environments, parasites are constantly attempting to penetrate host defences, which consist of multiple mechanisms to fight infections^3^. Such mechanisms include behavioural strategies (e.g. avoidance)^4^, the active reduction or elimination of pathogens (i.e. resistance) and/or the maintenance of homeostasis and fitness without interfering with pathogen burden (i.e. disease tolerance)^5–8^. While many studies in the past decades have explored the mechanistic and genetic bases of these different host defence strategies in diverse organisms ^9–11^, few have focused on how these processes can be shaped by evolution^12–15^. Experimental evolution is a powerful method by which it is possible to infer the evolutionary response of a host species to a specific pathogenic pressure under controlled conditions^16^. Previous studies of experimental evolution have addressed the impact of different hosts on parasite fitness^17^, of different pathogens on the host’s evolutionary trajectory^13,18,19^, of immune priming^20,21^, and of different routes of infection on host adaptation^19,22^.

It is postulated that oral exposure to pathogens constitutes a common process of infection in *D. melanogaster*^23,24^, as it lives on decaying substrates. When feeding on pathogenic microorganisms, *Drosophila* deploys different responses that include avoidance behaviours^25,26^, the barrier action of the peritrophic matrix^27^, localized high acidity of the midgut^28^, intensification of gut transit^29^, local production of Anti-Microbial Peptides (AMPs)^30^ and Reactive Oxygen Species (ROS) by gut epithelial cells^31,32^ and increase of enterocyte delamination and renewal rates^33–36^. In addition, *Drosophila* can activate a systemic humoral response, when the pathogen manages to cross the gut epithelium into the haemolymph^37,38^ or otherwise provokes a systemic reaction^23,39–41^. In the specific case of oral infection with the natural entomopathogen *Pseudomonas entomophila*, its high pathogenicity to *D. melanogaster* is linked to the production of toxins and virulence factors^42,43^ causing translational arrest in host tissues and the inability to renew damaged gut epithelium^44,45^. The above-presented plethora of possible host responses provides a clear picture of the multi-layered and mechanistically complex nature of the immune response, understood in its broad sense^3^.

In this work, we used a previously experimentally evolved outbred population of *D. melanogaster* (hereafter called “BactOral”) showing a fast evolutionary response to a strong selection against oral *P. entomophila* infection^19,46^, to investigate the mechanisms targeted by adaptation. We systematically tested for behaviour, resistance, and disease tolerance contributions, to provide a comprehensive account of the partitioning of these defense strategies in adaptation.

## RESULTS

### Evolved population shows higher survival upon infection

In previous work^19,46^, experimental evolution was conducted on an outbred population of *D. melanogaster* against oral infection either with the bacterium *P. entomophila* (hereafter, BactOral) or with a control solution (hereafter, Control). Since then, these populations^46^ have been kept under relaxed selection for over 80 generations. To understand to which extent this relaxation affected the response to the original selection regime, we measured survival of Control and BactOral populations upon *P. entomophila* oral infection. We found significantly higher survival of BactOral for both sexes (Fig. 1, Infected Control females *versus* Infected BactOral females: *z.ratio*= 9.123, *p*< 0.0001; Infected Control males *versus* Infected BactOral males: *z.ratio*= 9.909, *p*< 0.0001)(Suppl. Table 1). Moreover, these differences are not due to an inherent frailty of the Control as compared to BactOral. Indeed, survival under uninfected conditions for both long term longevity experiments (Suppl. Fig. 1 and Suppl. Table 2) and over the equivalent time period shows similar trends in both populations (Fig. 1, Uninfected Control females *versus* Uninfected BactOral females: *z.ratio*= 0.004, *p*= 1; Uninfected Control males *versus* Uninfected BactOral males: *z.ratio*= -1.080, *p*= 0.9611)(Suppl. Table 1).

**Figure 1.**
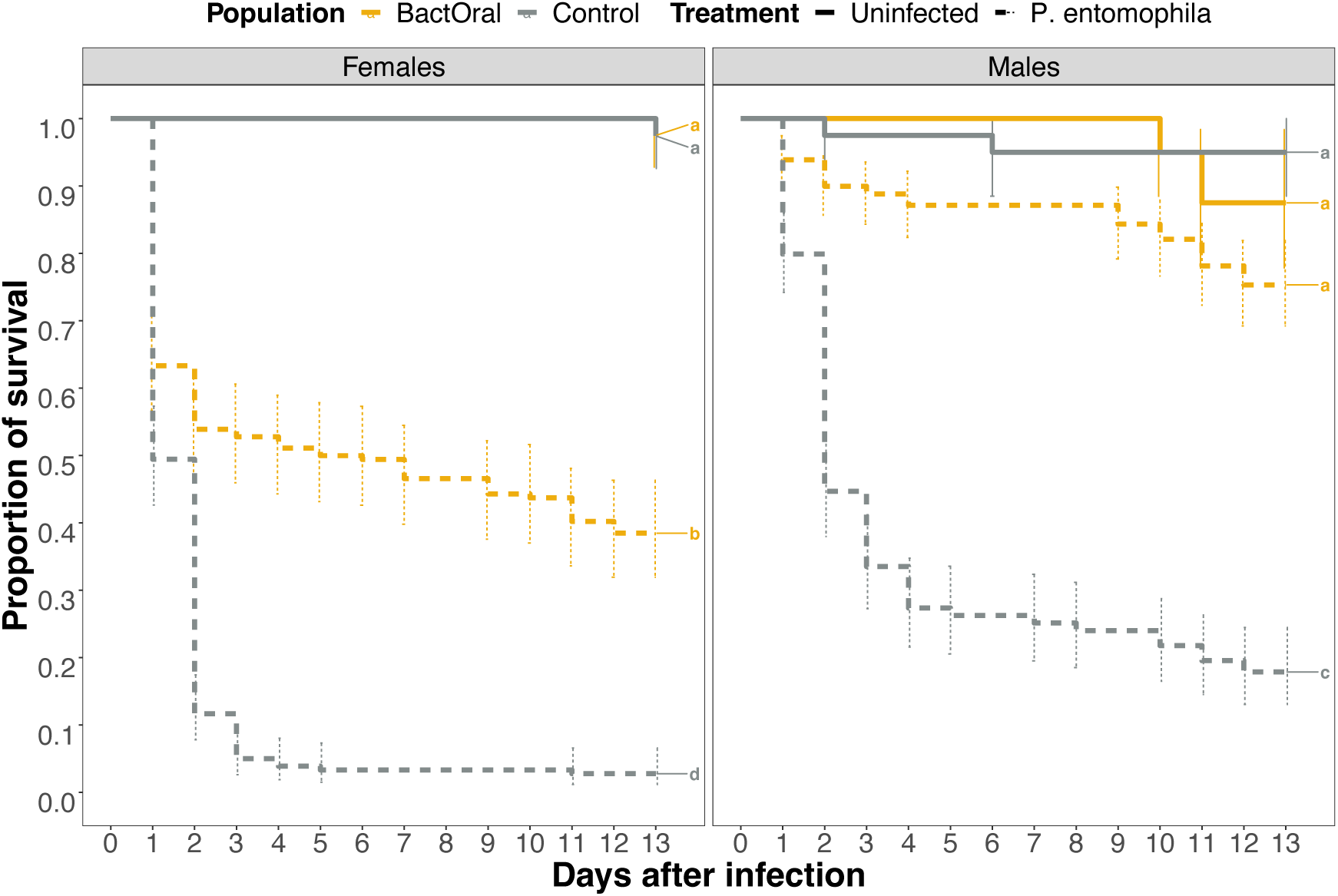
Survival upon infection with *P. entomophila*. Females (left) and males (right) of Control (grey) and BactOral (yellow) populations were orally infected with a culture of *P. entomophila* OD600=50 (dashed lines) or a control solution (solid lines) for 24 hours, after which they were flipped onto clean food and their survival measured daily. We identify a significantly higher survival after infection in BactOral (in both sexes) as compared with Control, in a treatment-dependent manner. Results from multiple comparisons are shown as letters in the plots. Sample sizes were 40 uninfected and 180 infected individuals, for each sex per population.

Taking these results into account, we undertook a systematic description of multiple traits previously associated to immune response to oral infections aiming to identify different processes targeted by selection and estimate their relative contribution.

### Evolved population has not changed bacterial uptake or defecation rates

Our systematic approach started by screening behavioural traits that impact bacterial numbers in the gut: feeding and defecation. To test for differences in feeding behaviour, we performed a Capillary Feeding assay (CAFE)^47^ on females of both Control and BactOral populations at different timepoints over 24 hours of exposure to bacterial suspension. We allowed individual flies to feed on a *P. entomophila* suspension mixed with food coloring in graduated glass capillaries and measured the amounts eaten by each female at 2, 8 and 24 hours of exposure. We found no significant differences between the amount ingested by individuals of the two populations (power analysis for a t-test: *Cohen’s d*= 1,06). Pairwise comparisons of their cumulative bacterial intake after 2 hours, 8 hours and 24 hours exhibited a comparable amount of total bacteria ingested (Suppl. Table 3 and Fig. 2A, Control *versus* BactOral at 2 hours, *t*= 0.097, *p*= 1; at 8 hours, *t*= 0.643, *p*= 0.9873; at 24 hours, *t*= 0.574, *p*= 0.9925).

**Figure 2.**
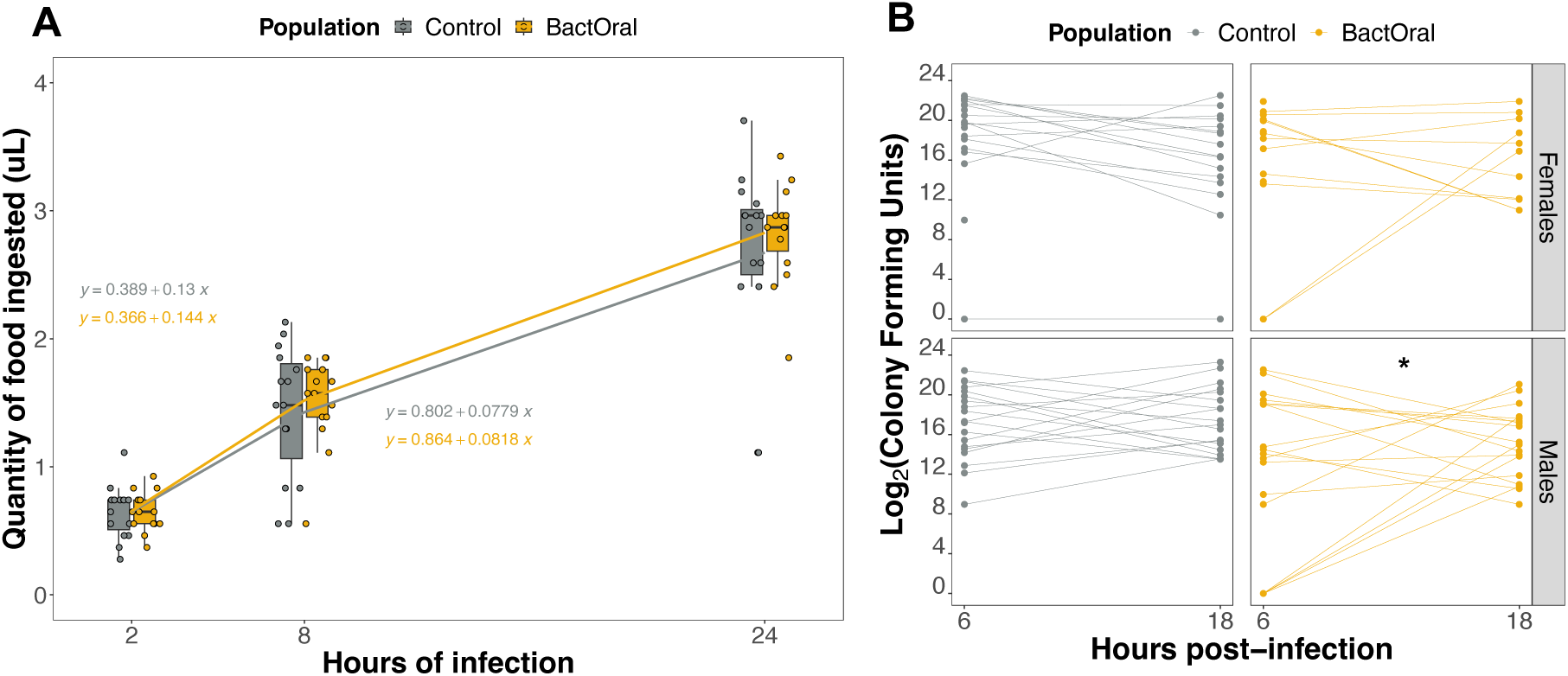
Evolved and control populations eat and defecate equivalent amounts of bacteria. A) CAFE assay. Females of Control and BactOral populations were fed individually *P. entomophila* solution through a graduated glass capillary and the amount of bacterial solution ingested (in µL) per fly was registered at 2, 8 and 24 hours. Cumulative number of µL ingested per fly showed no statistically significant difference in amounts ingested by either population at the different time-points. Linear regressions between averaged values eaten by each population between time-points show a comparable decrease in feeding as infection progresses (equations shown in plot for the periods, 2–8 hours (above lines) and 8–24 hours (below lines). 15 females were used from each population. **B)** Quantification of defecated bacteria. Females (top) and males (bottom) from Control (grey) and BactOral (yellow) populations were fed *P. entomophila* for 3 hours, after which the number *P. entomophila* defecated was estimated after 6 hours and 12 hours. Lines connecting the two points provide a sense of individual rates of live bacteria defecation within this time window. Linear regressions between the two counts of the same fly show a significant decrease in the number of bacteria defecated during the initial 6 hours and the following 12 hours by males of BactOral (shown with *). Additionally, there is a significant difference in the number of BactOral flies that defecated below our detection threshold (zero CFU counts) at 6 hours compared to 18 hours and compared to Control for both timepoints. Samples sizes are shown at the top of each panel.

Additionally, we estimated the average slopes from linear regression where microliters of bacteria eaten were modelled between different time points, concluding that the bacterial quantities ingested by both populations remain undistinguishable as the infection progresses. Thus, exposure affects feeding ability of both populations to a comparable degree and, consequently, adaptation to oral infection with *P. entomophila* in the BactOral regime did not involve an alteration of feeding behaviour.

Considering the absence of significant differences in bacterial intake, an alternative would be that the BactOral population had evolved a more effective bacterial defecation, which would reduce the time of pathogen permanence and proliferation in the gut. We established a protocol to measure live bacteria loads in the faeces, hence considering the number of bacteria defecated as a proxy for defecation rate itself, assuming that no interaction between the two parameters exists. To this purpose, we took individual flies upon oral infection and quantified the CFUs defecated over a six-hour and the subsequent twelve hour-periods, post-exposure (Fig. 2B).

We observed a few instances where flies did not expel alive bacteria during that period or did so in a quantity that fell below our detection threshold, resulting in a few counts of zero. Therefore, a zero-inflated model with a negative binomial distribution was fitted on the counts data (accounting for overdispersion) jointly with a binomial distribution on the zeros. For the counts portion of the model, none of the three factors significantly predicted the measured bacterial loads (Suppl. Table 4, model parameters - Population: *z.value*= 1.324, *p*= 0.185; Sex: *z.value*= 0.073, *p*= 0.942; Timepoint: *z.value*= -0.359, *p*= 0.720). However, looking at the zero portion of the model, we find a significant effect of Population (Population: *z.value*= - 2.407, *p*= 0.016; Timepoint: *z.value*= -0.006, *p*= 0.996).

We then performed post-hoc pairwise comparisons on these regressions and identified a significant decrease in bacterial loads defecated by males of BactOral, between the two timepoints (males: *z.ratio*= -2.719, *p*= 0.033), but no other significant effects among bacterial counts between populations and/or timepoints, in a sex-dependent manner. However, when considering the zeros, we found a significant difference in the proportion of BactOral flies with undetectable amounts of bacteria in the faeces between the first and second timepoints (B6h– B18h: *z.ratio*= 4.092, *p*= 0.0002), as well as compared to Control at both timepoints (B6h– C6h: *z.ratio*= 3.612, *p*= 0.002; B6h– C18h: *z.ratio*= 3.530, *p*= 0.002)(Suppl. Table 4).

These results demonstrate that both Control and BactOral populations defecate alive bacteria in comparable amounts upon infection. Nonetheless, we detected that males of BactOral decrease the amount of live bacteria they defecate as time progresses, hinting at a possible resistance mechanism more evident in males of the adapted lines, that was not significant in females. This observation is corroborated by the difference in the number of “zeros” found between both populations, with BactOral exhibiting more instances of expelling undetectable bacterial loads.

### Evolved population shows a faster decrease and clearance of bacteria

Next, we studied the physiological responses to estimate the roles of resistance and disease tolerance. First, we tested for structural changes in the gut itself that could influence the “carrying capacity” of the structure and create difficulties in interpreting the quantification experiments. We measured the capacity of BactOral to regenerate its gut epithelium upon bacterial infection relative to Control, using initial length and extent of the shortening induced by infection^36^, as a proxy. For this, we measured the total length of the guts of females, before and after 24 hours of oral infection with *P. entomophila* and performed multiple pairwise comparisons on a linear model with mixed effects to pinpoint how population and infection status affected this trait. We found a shortening of approximately 2 mm in the total length of infected guts by comparing the post-infection timepoint of 24 hours to the initial timepoint, prior to exposure, in both populations (Suppl. Table 5 and Suppl. Fig. 2, Control before infection *versus* Control 24 hours post-infection: *t*= 6.274, *p*<0.0001; BactOral before infection *versus* BactOral 24 hours post-infection: *t* = 5.453, *p* <0.0001). Nonetheless, no significant differences between populations in the total length of the guts could be detected both before and after infection (Without infection: BactOral *versus* Control – *p*= 0.6719; after 24 hours of infection – *p*= 0.9925), (power analysis for a multiple linear regression: *Cohen’s d* for Population= 0,71; *d* for Treatment= 0,71) (Suppl. Fig. 2).

To explore the dynamics of bacterial proliferation inside the host, we performed a time-course characterization of bacterial loads during and after oral infection by counting the number of colony-forming units (CFUs) obtained from individual flies. In Figure 3, we show the progression of bacterial loads from 1 hour until 52 hours post-infection (Fig. 3A) as well as the cumulative number of individuals without detectable live bacteria (Fig. 3B). The infection dynamics of both populations start with very high numbers of *P. entomophila* at early stages, confirming the absence of differences in initial inoculum reported above (Table 1, parameter *a*: 3.072± 0.016). Bacterial loads continuously decrease and, at approximately 20 hours of infection, Control and BactOral population begin to differ significantly in a sex-dependent manner. From this point onwards we observe a steady decrease of bacterial quantities in the two populations, always more severe in males than in females, and in BactOral than in Control flies (Fig. 3A). In addition, the number of flies that clear the infection also increases with time in both populations, but at a faster rate and reaching higher levels in BactOral (Fig. 3B). By 52 hours post-infection, in both males and females, the number of “cleared” flies in the BactOral population was twice as high as that of its Control counterpart (Fig. 3B). Together, these analyses indicate that the evolved population eliminates bacteria faster and more efficiently than Control.

**Figure 3.**
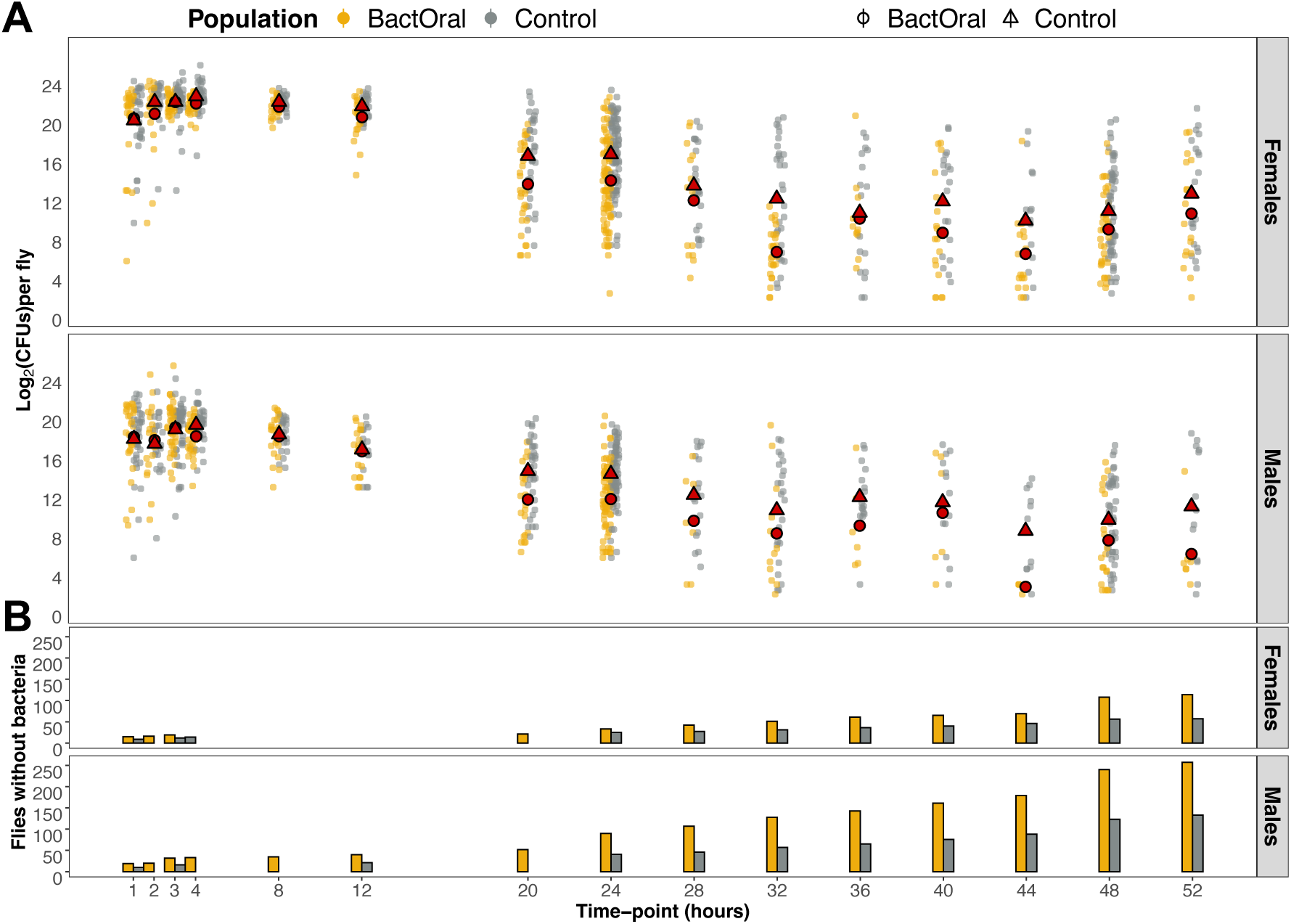
Bacterial load dynamics. Individual flies from Control (grey) and BactOral (yellow) were collected upon exposure to *P. entomophila* at different timepoints for assessment of the number of Colony Forming Units (CFUs) inside their bodies. After the end of the infection period (24 hours), flies were changed onto clean food and collection continued until 52 hours post-infection. **A)** Log2 of the number of CFUs found for females (top) and males (bottom) of either Control (grey) or BactOral (yellow) populations, with the mean value at each time-point evidenced in red (triangles for Control and circles for BactOral). There are no detectable differences in bacterial loads between both populations until 20 hours of infection, when BactOral starts to evidence a bigger decrease in bacterial counts. **B)** Cumulative number of individual flies from which no CFUs were obtained increases as time progresses, with the BactOral reaching significantly higher final values. Numbers at the top of each panel correspond to the sample size at each timepoint, with BactOral population in yellow and Control population in grey.

**Table 1.**
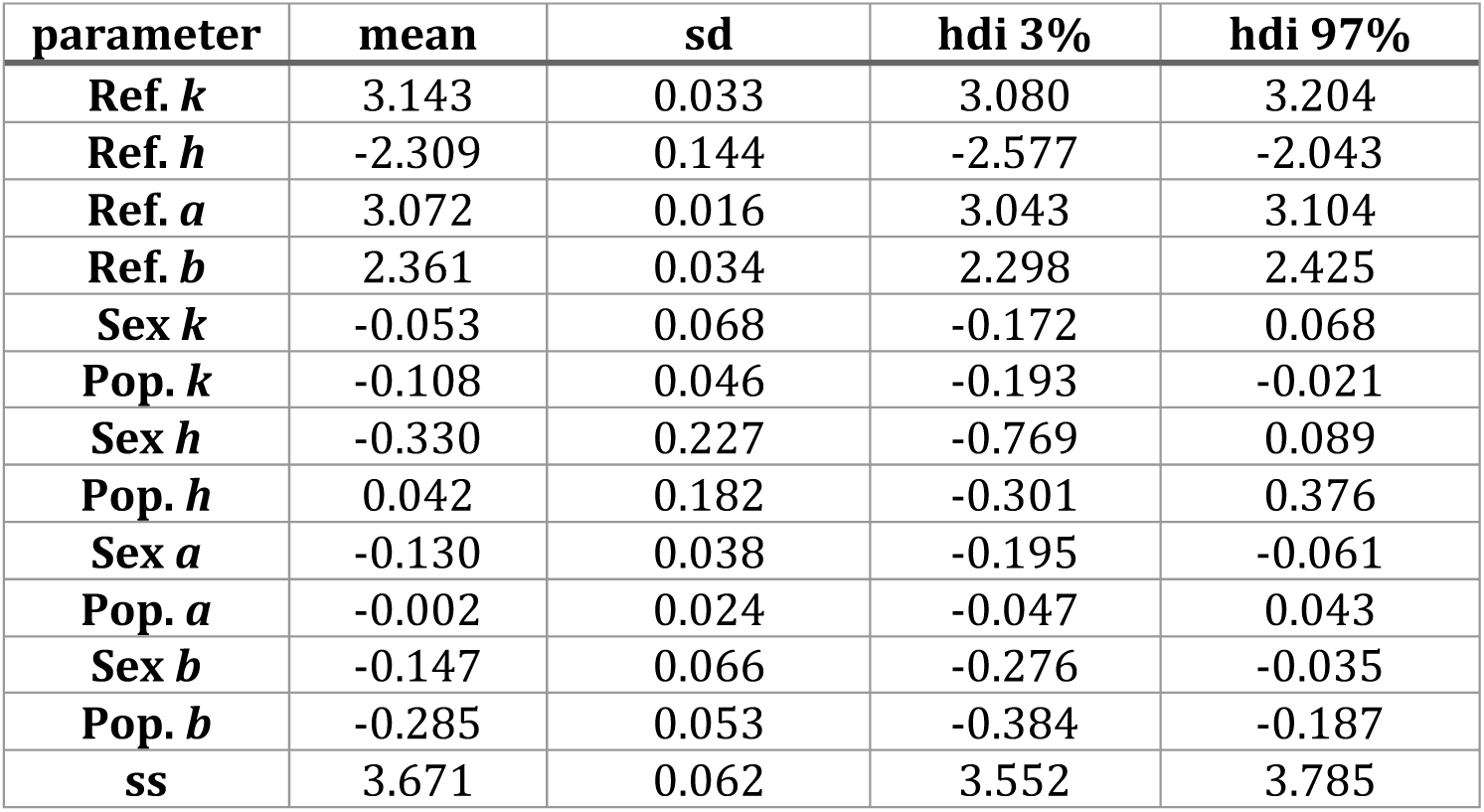
Parameter estimates for the model sigmoidal function fitted to the time- course of bacterial counts. A sigmoidal function of 4 parameters (*a*, *k*, *h*, *b*) was fitted to the bacterial dynamics time-course shown in Figure 3A, for quantification of differences between Control and BactOral CFU counts. Control females were used as reference on which to estimate the contribution of tested variables (variable “Sex”: males and variable “Population”: BactOral) on the parameters. Effects (Sex and Population) on parameters were considered significant when the 94% highest posterior density interval (hdi) did not include zero. The “ss” parameter represents the estimate of the standard deviation of the normal error structure.

| parameter | mean | sd | hdi 3% | hdi 97% |
| --- | --- | --- | --- | --- |
| **Ref. *k*** | 3.143 | 0.033 | 3.080 | 3.204 |
| **Ref. *h*** | -2.309 | 0.144 | -2.577 | -2.043 |
| **Ref. *a*** | 3.072 | 0.016 | 3.043 | 3.104 |
| **Ref. *b*** | 2.361 | 0.034 | 2.298 | 2.425 |
| **Sex *k*** | -0.053 | 0.068 | -0.172 | 0.068 |
| **Pop. *k*** | -0.108 | 0.046 | -0.193 | -0.021 |
| **Sex *h*** | -0.330 | 0.227 | -0.769 | 0.089 |
| **Pop. *h*** | 0.042 | 0.182 | -0.301 | 0.376 |
| **Sex *a*** | -0.130 | 0.038 | -0.195 | -0.061 |
| **Pop. *a*** | -0.002 | 0.024 | -0.047 | 0.043 |
| **Sex *b*** | -0.147 | 0.066 | -0.276 | -0.035 |
| **Pop. *b*** | -0.285 | 0.053 | -0.384 | -0.187 |
| **ss** | 3.671 | 0.062 | 3.552 | 3.785 |

In order to pinpoint more accurately the parameters contributing to this overall difference, we modelled the bacterial load and zero dynamics, and assessed differences in parameter estimates considering the initial load or inoculum (*a*), the load in the long-term or set-point bacterial load (*b*), the timepoint at which the load is reduced in half (*k*) and the highest rate of decrease or slope at timepoint *k* (*h*) (see Materials and Methods for details). In Table 1, we show the mean estimates, standard deviation, and high-density intervals (3% and 97%) for all four parameters of the fitted sigmoidal model, for Control and BactOral populations. Overall, these results show that sex has a significant effect on all curve parameters (*k*= -0.053 ± 0.068; *h*= -0.330 ± 0.227; *a*= -0.130 ± 0.038; *b* = -0.147 ± 0.066), reinforcing the notion of a strong sexual dimorphism in response to infections^48–50^. Additionally, populations differ on “time to half height” (*k*= -0.108 ± 0.046) and final load (*b*= -0.285 ± 0.053), but not on initial load (*a*= -0.002 ± 0.024) nor on highest rate of decrease (*h*= 0.042 ± 0.182).

Because this time-course experiment yielded a significant amount of zero counts, corresponding to individual flies for which no CFUs could be counted, we treated this as an independent dataset for which we developed a separate model focused on infection clearance (Table 2). With that, we found that *a1* (final fraction of zeros) was significantly affected by both sex and population (with no observable overlap between the mean estimates for each condition) (*a1* Females= -0.781 ± 0.07; *a1* Males= -0.061 ± 0.069; *a1* Control= -0.700 ± 0.070; *a1* BactOral= -0.144 ± 0.069), with more BactOral individuals having cleared the infection by the end of the experiment, and males more so than females. There were significant overlaps between the estimates of the remaining parameters, evidencing a difference in clearance rate only in late stages of bacterial infection. In sum, these data align with those of bacterial load dynamics, whereby flies from BactOral population clear *P. entomophila* faster and better from 20h of exposure onwards.

**Table 2.**
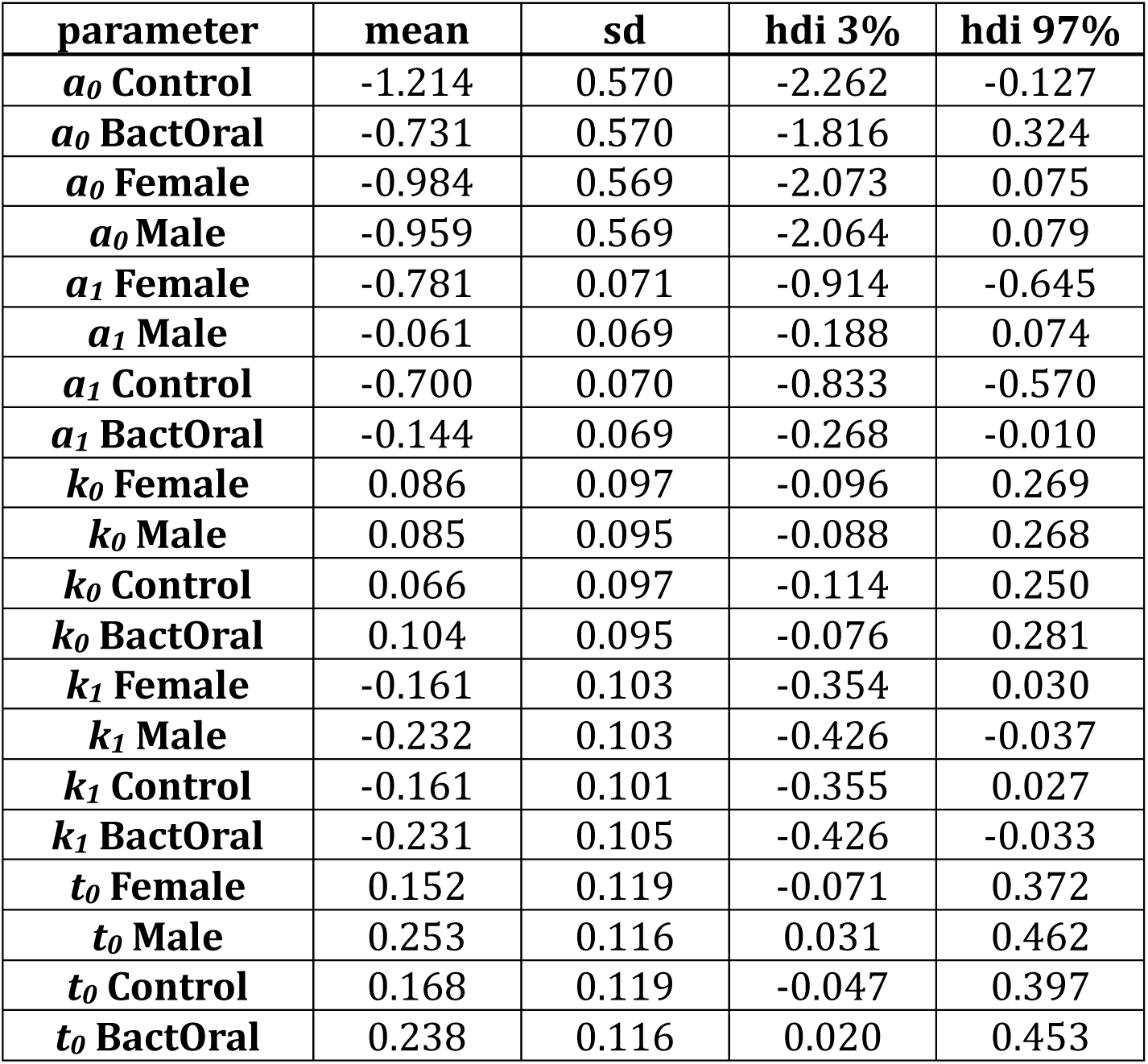
Parameter estimates for the model logistic function fitted to the zero counts of the time-course of bacterial counts. An adapted logistic function of 4 parameters (*a0*, *a1*, *k0*, *k1*) was fitted to the progression of zero counts shown in Fig. 3B), for quantification of bacterial clearance differences between Control and BactOral. Single estimates were generated for the contribution of each variable (Sex: females and males; Population: Control and BactOral) to the model parameters. Effects were considered significant when was considered when the mean estimate ± standard deviation did not overlap for the same parameter between each variable, which is the case for *a1* when considering Population and Sex. High-density intervals of 3% and 97% show that all our estimated parameters have a high-density probability (i.e. credibility) as they are all included in them.

These results support the hypothesis that resistance mechanisms participate in the adapted response of BactOral. Indeed, bacterial numbers are affected differently between BactOral and Control populations despite the equivalent initial bacterial inocula (Fig. 2A, Table 1) and rates of live bacteria defecation (Fig. 2B). The differences in number of detected live bacteria inside flies suggests a direct bacterial control mechanism is operating more efficiently in BactOral individuals.

### Differences in AMP expression are consistent with distinct bacterial loads dynamics

In an attempt to pinpoint which resistance mechanisms contribute to the differences in bacterial clearance rates observed between populations, we infected female flies from Control and BactOral and dissected their guts prior to exposure, and after 8, 24, and 32 hours of infection with *P. entomophila*. We quantified expression levels for a panel of 10 AMPs covering different molecular families, dedicated pathways and pathogen specificities, including *Attacin-A*, *Bomanins Short 2* and *3*, *Cecropin-A1*, *Defensin*, *Diptericin-A*, *Drosomycin* and *Drosomycin*-like *2*, *3* and *5* (Fig. 4, Suppl. Tables 6-7, and Supp. Fig. 3).

**Figure 4.**
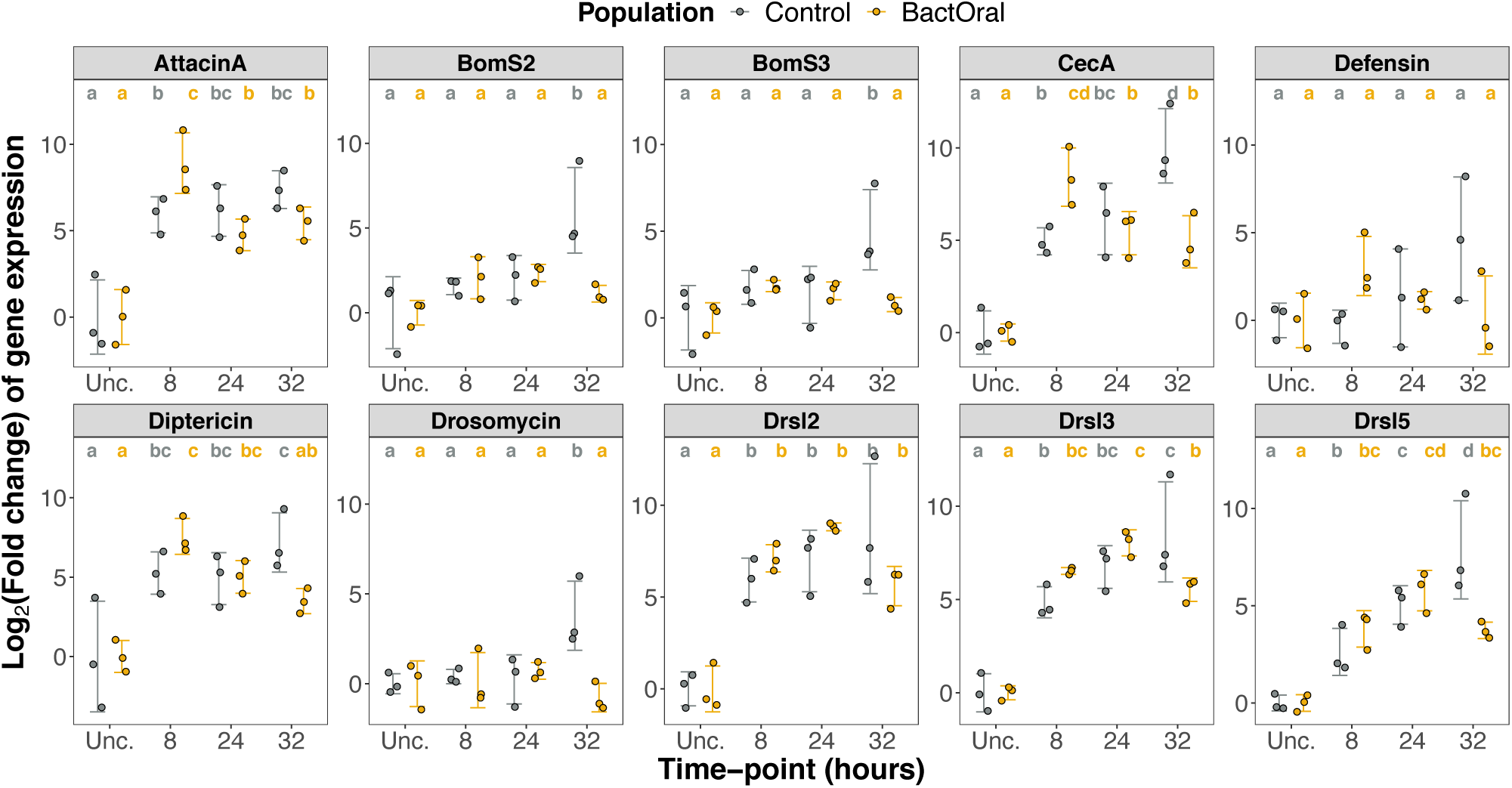
AMP expression differs between populations. Guts from Control (grey) and BactOral (yellow) female flies (3 samples containing 10 guts, each) were dissected before exposure (“Unc.”) and at different timepoints after oral infection (8, 24 and 32 hours) with *P. entomophila.* Relative gene expression, normalized to housekeeping gene *EIF2,* was performed for a representative panel of 10 AMPs. We detected a significantly higher up-regulation in BactOral compared to Control for *AttacinA* and *CecropinA* at 8-hours of infection. Results from these comparisons are shown in the letters above the plots.

We first verified if the baseline levels of AMPs (i.e., expressed under homeostatic conditions) differed between populations, by relativizing expression levels to the Control population at the unchallenged timepoint. With this approach, we established that BactOral and Control populations express comparable amounts of 8 AMPs when unchallenged, the two exceptions being *Cecropin-A1* (*CecA*) with lower expression levels (*t* = 2.892, *p*= 0.0445) and *Drosomycin-like 5* (*Drsl5*) with higher levels (*t* = -4.686, *p*= 0.0094) (Supp. Fig. 2) in the Control compared to the BactOral population.

In addition, and since our main goal was to characterize the temporal dynamics of AMP expression in both populations, we performed the analysis for the entirety of the time-course using the unchallenged timepoint as reference (power analysis for a t-test: *Cohen’s d*= 0,62) (Fig. 4 and Suppl. Tables 6-7). Overall, we detect consistent dynamics with the reduction of expression happening sooner (by 32 hours) in BactOral than Control across 7 of the 10 AMPs tested, the exceptions being *Attacin A*, *Defensin* and *Drosomycin-like 2* (*Drsl2*) (Fig. 4). This earlier down-regulation in BactOral relative to Control flies (at 32-hours) of *BomaninS2* (*BomS2*) (*t* = 4.347, *p*= 0.0047), *BomaninS3* (*BomS3*) (*t* = 4.035, *p*= 0.0089), *CecA* (*t* = 4.529, *p* = 0.0010), *Diptericin* (*t* = 2.613, *p*= 0.0439), *Drosomycin* (*t* = 4.795, *p*= 0.0056), *Drosomycin- like 3* (*Drsl3*) (*t* = 3.194, *p*= 0.0105) and *Drsl5* (*t* = 4.266, *p*= 0.0021). Furthermore, we detected a significantly higher up-regulation of *AttacinA* and *CecA* in BactOral at 8 hours of infection (*AttacinA*: Control 8h *versus* BactOral 8h – *t* = -2.563, *p*= 0.0389; *CecA*: Control 8h *versus* BactOral 8h – *t* = - 3.034, *p*= 0.0123), consistent with the role of these peptides as main players against infections with Gram-negative bacteria^51,52^. A similar trend was observed for other tested AMPs such as *Defensin* and *Drsl2*, although these changes were not statistically significant (*Defensin*: Control 8h *versus* BactOral 8h – *t* = -2.101, *p*= 0.231; *Drsl2*: Control 8h *versus* BactOral 8 h.p.i. – *t* = -0.899, *p*= 0.465).

Principal Component Analysis (PCA) supports these trends (Suppl. Fig. 4). The two main Principal Components (PC1 and PC2) evidence the time-dependent differences observed between populations by showing an initial indistinguishable grouping of both, followed by a dissimilar change in expression patterns at 8 hours of infection which culminates with a maximum divergence between Control and BactOral at the later timepoint (32 hours).

### Increase of AMP levels in the Control population phenocopies BactOral response to infection

Having correlated the improved bacterial clearance of BactOral flies (Fig. 3) with significant differences in expression levels of AMPs between populations (Fig. 4), we sought to functionally associate these two observations. For that, we conducted a priming experiment to induce higher basal levels of AMPs by feeding flies a solution of heat-killed bacteria. These primed flies were, subsequently, infected orally with live *P. entomophila*, and followed daily for mortality. This exposure to dead bacteria has been shown to “artificially” increase AMP levels^53–55^. We expect the priming with heat-killed bacteria to increase the basal expression of AMPs and thus provoke, particularly in Control flies, an up-regulation of the immune system, allowing test of the influence of this heightened condition in survival to oral infection.

Flies previously fed with heat-killed bacteria increased survival to infection with live *P. entomophila* in BactOral and Control populations, confirming the stronger immune response expected upon priming (Suppl. Table 8 and Fig. 5, BactOral females fed with live bacteria (control) *versus* BactOral females primed with heat-killed bacteria (primed): *z.ratio*= 7.193, *p*< 0.0001; BactOral males treated with control *versus* BactOral males primed with Heat-killed bacteria: *z.ratio*= 4.368, *p*= 0.0001; Control females fed with control *versus* Control females : *z.ratio*= 10.011, *p*< 0.0001; Control males treated with control *versus* Control males primed : *z.ratio*= 7.299, *p*< 0.0001). Females from the Control population primed with heat-killed bacteria increased their survival to oral infection, virtually phenocopying BactOral infected with live bacteria, whereas in males, the same tendency is observed but to a lesser degree (Suppl.

**Figure 5.**
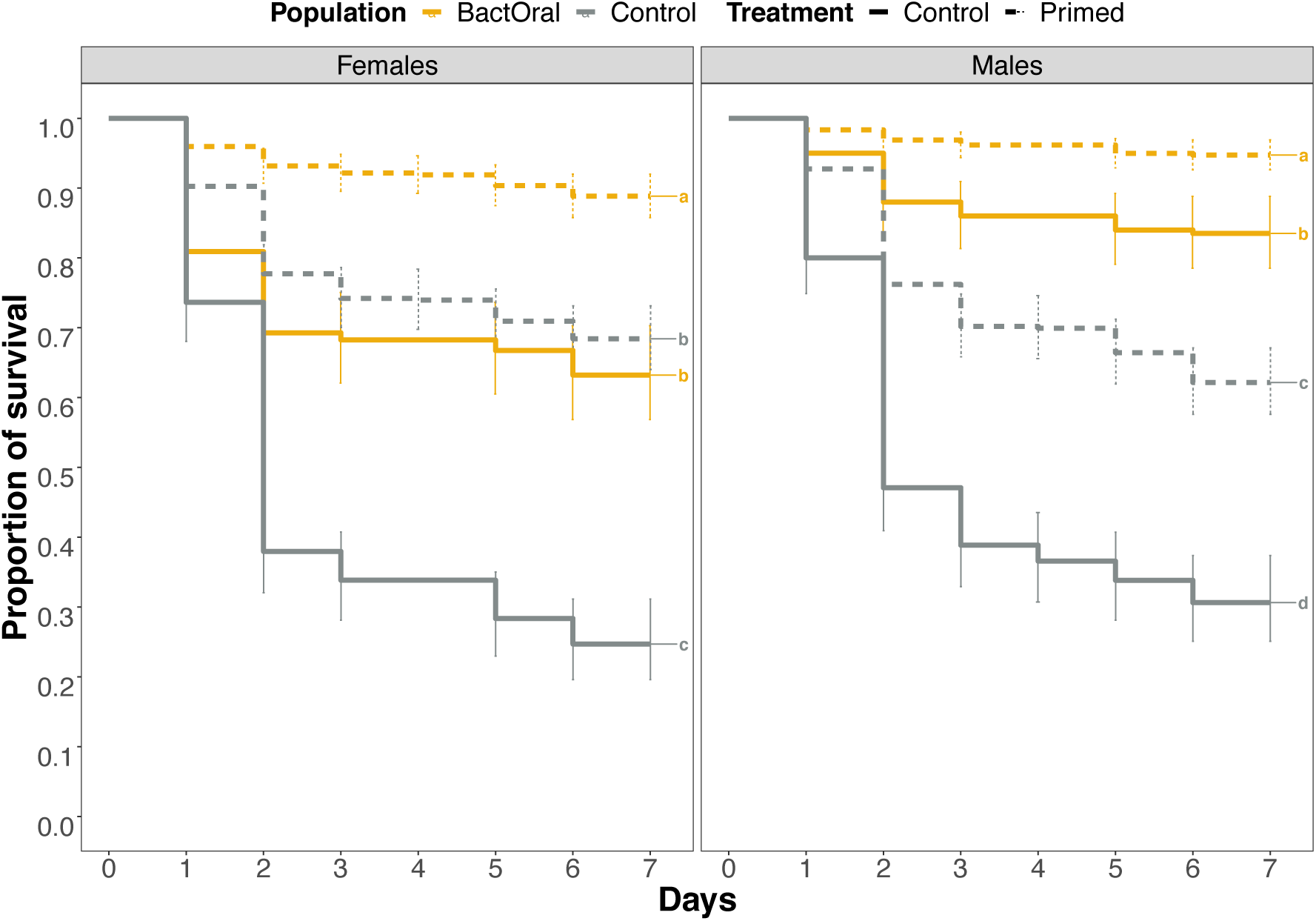
Control flies primed with heat-killed bacteria phenocopy BactOral survival to oral infection with *P. entomophila*. Control (grey) and BactOral (yellow) females (left) and males (right) were fed a solution of heat-killed bacteria for 2 hours before oral infection (dashed lines) or directly infected with live *P. entomophila* (full lines) for 24 hours, after which they were changed onto clean food and their survival measured daily. Previous exposure to heat-killed bacteria before infection increased survival in both populations, leading additionally Control females to phenocopy BactOral survival levels upon infection. Results from multiple comparisons are shown as letters in the plots. Numbers at the end of each trajectory correspond to the sample size, with BactOral population in yellow and Control population in grey.

Table 8 and Fig. 5, BactOral females fed with control *versus* Control females primed: *z.rati*o= 1.629, *p*= 0.3620 and BactOral males fed with control *versus* Control males primed: *z.ratio*= - 4.547, *p*<0001). This effect is supported by a time-course of relative expression levels of IMD pathway key players, *Relish* (its devoted transcription factor) and its associated AMP, *Diptericin*, as well as the Toll-activated AMP, *Drosomycin* (Suppl. Fig. 5). For both populations, we observed an increase in levels of *Diptericin* to a comparable degree upon exposure to heat-killed *P. entomophila*, but not of *Drosomycin*. With this experiment we strengthened the case for a role of improved resistance in the adapted phenotype of the BactOral population.

### BactOral maintains higher survival upon immune response activation

To ascertain whether disease tolerance may also contribute to BactOral increased survival upon oral infection, we continuously fed flies with a mixture of heat-killed *P. entomophila* and followed survival (Fig. 6).

**Figure 6.**
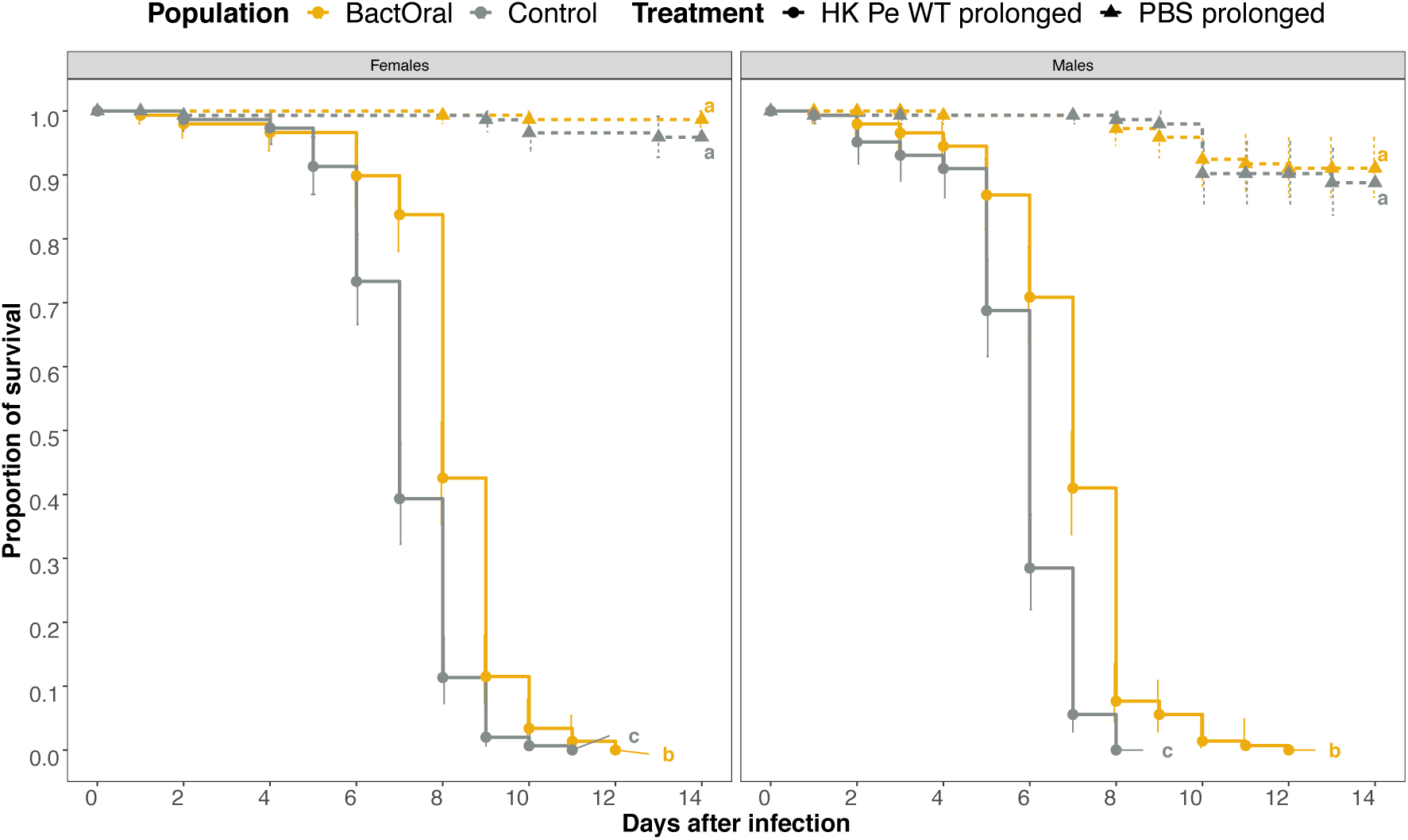
BactOral displays higher survival than Control upon immune system activation without bacterial proliferation. Daily survival was measured for Control (grey) and BactOral (yellow) populations fed a mixture of food with heat-killed bacteria or PBS continuously. BactOral females and males exhibit a slower mortality compared to Control flies until around day 8. This difference fades away for both sexes after that period. Survival comparisons between populations and treatments are shown as lower-case letters. Sample sizes were 150 individuals, for each sex, per population, per treatment.

With this setup we verified that both females and males of BactOral survive significantly more than Control after the specific damages induced by constant exposure to heat-killed bacteria. Importantly, the survival differences detected between populations occur up until the time at which flies were selected to reproduce during the experimental evolution protocol (10 days upon infection), evidencing the adaptive nature of this response. (Suppl. Table 9 and Fig 6, Control - HK *versus* BactOral - HK females: *z.ratio*= 6.579, *p*< 0.0001; males: *z.ratio*= 9.612, *p*< 0.0001). No significant differences exist under control conditions (Control - PBS *versus* BactOral - PBS females: *z.ratio*= 1.392, *p*= 0.504; males: *z.ratio*= 0.587, *p*= 0.936).

## DISCUSSION

In this work, we enquired systematically for the bases of adaptation of one evolved line of *D. melanogaster*, against an oral bacterial infection considering the three general layers that constitute the immune response: behaviour, resistance and disease tolerance^3,8,56^. Upon experimental evolution of an outbred population of *D. melanogaster* against oral infection with *P. entomophila*, and after approximately 6 generations, the selected population (BactOral) increased survival upon infection^19^. This phenotype was maintained throughout the selection experiment but, since then, the population has been kept for more than 80 generations under relaxed selection. Despite the fact that no physiological costs associated to this response could be detected in any of the original four replicates^46^, we tested the immunocompetence of one selected representative replicate per evolved population. The average survival rate after oral infection of flies from BactOral was approximately 63% (50% for females and 75% for males) and approximately 10% in the Control population (5% for females and 15% for males). Considering that at the end of selection, BactOral survival reached 90 %^19^, this partial decrease in overall survival after infection with bacteria can potentially underestimate the contributions of each of the different mechanisms behind adaptation. Importantly, these losses could be attributed to genetic drift acting on the populations during the long relaxed-selection period they sustained. Notwithstanding, we established that it is methodologically possible to ascertain the relative importance of behavioural, resistance, and disease tolerance mechanisms.

### No role for behavioural or structural differences

It is well described that the route of infection is of paramount importance to the outcome of host-parasite interactions. In *D. melanogaster*, the mode of entry of a bacterial pathogen into the host’s body elicits different physiological responses^19,57–61^, which can start by the avoidance of the pathogenic substrate, in the case of infection through oral route^26,62,63^, both by inhibiting feeding or choosing oviposition sites according to bacterial composition^26,53,62–64^.

*P. entomophila* is a natural pathogen of Drosophila that causes high mortality when ingested across diverse genetic backgrounds and conditions^23,43,44,65,66^. Thus, it is plausible to consider that fasting could constitute a general response against oral infection. The simplest behaviour that could explain the observed differences in survival between populations would be that BactOral flies evolved a reduced intake during exposure to bacteria, thereby reducing their “inoculum” size. Indeed, fasting is a prevalent response to infection and behavioural avoidance is a common strategy to limit exposure to infected substrates in different systems^4,62,63^. However, we showed similar amounts of bacteria were ingested by females of both populations (Figure 2A)^23,67^. Moreover, we consubstantiate the findings from the CAFE assay with the bacterial loads time-course, that show equivalent amounts of *P. entomophila* colonies inside infected flies at early timepoints. Our results indicate that behavioural inhibition of bacterial intake was not one of the mechanisms that evolved in BactOral flies, eliminating fasting as a component of the adaptation to *P. entomophila* exposure.

Previous reports have shown that pathogen intake can affect gut function and structure^34–36,45^ as well as influence the dynamics of exposure by altering defecation rates^29^. However, we could not find evidence for any differences in these features between control and evolved populations. Firstly, there are no changes in gut length in steady-state conditions that would somehow enable the BactOral population to *a priori* face bacterial infection more favourably. Increased enteric cell renewal, which often leads to a temporary shortening of gut length^36^, has been described as one of the main mechanisms of *D. melanogaster* defense against oral bacterial infection^35,36,45^. In our experiments, we detected a significant decrease in total gut length after 24 hours of oral infection, but this change was equivalent in both populations, not supporting a role for altered renewal rate in BactOral adaptation.

These experiments, however, do not completely discard the hypothesis of differential gut structure underlying the adapted phenotype of BactOral. For example, proportions of gut cell types can change upon infection^68^ and such qualitative shift could contribute to higher fitness upon exposure to *P. entomophila* in the adapted population. Also, it could be that relevant changes in gut length occur at later timepoints. Notwithstanding, it has been previously shown that infection-induced gut cell loss starts as early as 2 hours after exposure, with a return to 75% of the total original gut length after 24 hours^36^. This time window is well within the boundaries defined in our setup. Another important aspect to note is that renewal capacity is sexually dimorphic under diverse physiological contexts^69,70^. For instance, females have higher gut cell renewal rates than males upon oral infection with *P. carotovorum* 15 ^71^, a bacterium that provokes higher mortality in males than females, contrary to oral infection with *P. entomophila* (see Results). The disparity between those and our observations may be explained by the different mechanisms of virulence employed by these two pathogens to infect *D. melanogaster*^45^. Be as it may, we envision further experiments testing for potential differences in gut delamination rates between evolved populations at distinct timepoints.

BactOral flies also did not expel pathogenic bacteria from their guts differently from Control. However, males of BactOral show a decrease in the number of alive bacteria defecated between 6- and 18-hours post-exposure, evidencing, to some extent, improved bacterial clearance and suggesting a role for resistance. Since our protocol only detects live bacteria, we could potentially be missing an important component of the response and future work should quantify defecation of alive and dead bacteria, for example through fluorescent microscopy, to compare populations with regards to the contribution of overall peristalsis to the elimination of total gut content^29^. In addition, considering the slightly higher number of of BactOral flies that defecated alive bacteria below our detection limit (zeros), we hypothesize that this population either retains *P. entomophila* for longer inside guts before they begin to expel it (which could point to higher disease tolerance), or reduce bacterial loads more effectively at earlier stages, even though, with time, bacterial proliferation recovers (again suggesting stronger resistance).

### A role for resistance

To further test the suggested role for resistance, we characterized the bacterial load dynamics in evolved populations over 52 hours and determined that BactOral flies eliminate pathogens in a more efficient way than Control. This significant change in bacterial loads indicates that BactOral has evolved higher resistance, that control earlier and clear better gut bacterial loads. As this model was applied to the logarithm of mean bacterial counts at each timepoint, it is possible that information regarding intra-populational variation is not considered in the analysis. Nevertheless, the conclusions from the comparison of the bacterial counts are further supported by the separate analysis on the number of “zeros” as a function of time, which is a proxy for bacterial clearance.

Even though there is no discernible difference in the amount of CFUs infecting both males and females between both populations until approximately 20h of infection, by this time individuals from BactOral begin to exhibit gradually decreasing levels of bacteria. This trend aligns with the higher clearance rate observed in the adapted population (and even higher in males than females) as measured by the increase in the number of flies from which we cannot grow colonies of *P. entomophila* (“zeros”). Flies from the Control population control their bacterial loads (at around 20h of infection) but do it at a slower pace and in a less effective way (i.e. many individuals exhibited high bacterial counts by the end of the experiment). It is possible that our proxy for clearance (i.e., flies classified as “zeros”), could be overestimated due to a low detection threshold. Indeed, it has been shown that *D. melanogaster* can endure chronic bacterial infections^72–74^, albeit at rather low levels, which can be the case in our experimental setup. Nonetheless, this holds true for both populations, therefore not invalidating our interpretation of a different bacterial clearance rate between genotypes.

Having shown that BactOral clears *P. entomophila* infection more rapidly, we hypothesized that this response could rely on an increased production of immune effectors, such as AMPs. Analysis of AMP gut gene expression showed that in unchallenged condition, only two AMPs (*CecA1* and *Drsl5*) differed in expression between populations (down- and up-regulation respectively), indicating that the base levels of AMP expression are unlikely to have been a major target of selection. The detected expression difference of *Drsl5*, a gut-specific AMP peptide, could represent a pre-emptive resistance mechanism acquired by BactOral, by establishing a more antagonistic environment for *P. entomophila* proliferation in the guts of these flies after ingestion.

At 8 hours of exposure, comparing BactOral to Control, two AMPs show a significantly faster up-regulation while seven of the panel members display a pattern of faster downregulation, later on (32h). These results suggest that a two-step process might be taking place, whereby the response is initially (8h) more aggressive, permitting a more effective bacterial control, and is turned off sooner (32h), limiting the secondary effects of prolonged immune responses (immunopathology). Interestingly, *Drosomycin-like* genes (*Drsl2*, *Drsl3*, and *Drsl5*), whose expression is specific to the gut epithelium and are regulated by JAK- STAT^35^, show up-regulation after 8-hours of infection in both populations. For all three genes, there is a slight tendency for a faster response that could represent a more effective bacterial control in the BactOral population and a faster down-regulation at the 32-hour timepoint that could limit self-inflicted damage. Likewise, we observed a similar process of down-regulation of other AMPs specifically deployed to fight off infections with Gram-negative pathogens (e.g. *CecA1* and *Diptericin*) at the 32-hour timepoint. In contrast, some AMPs typically associated to response to Gram-positive bacteria, (e.g. *BomS2*, *BomS3* and *Drosomycin*^51,75,76)^, do not exhibit significant up-regulation after exposure to *P. entomophila* until 32-hours post-infection and only in the Control population.

Strikingly, and even though *Diptericin* has been shown to be deployed by *D. melanogaster* to resist *P. entomophila* infection^67^, we did not detect a difference in the relative expression of this gene between Control and BactOral until after 24 hours of infection. At this timepoint, adapted flies are already returning to basal levels of AMP production whereas Control flies keep actively fighting off the infection.

The same pattern is observed also in the priming experiment (Fig. 5 and Suppl. Fig. 5). After priming with heat-killed bacteria and subsequent exposure to live bacteria, both populations maintain an equivalent expression increase until 8 hours but from 24 hours onwards BactOral shows, again, an earlier downregulation of *Diptericin* and a quicker return to homeostatic levels of expression (Suppl. Fig. 5). In brief, the dynamics of *Diptericin* expression mimic the expected effect of priming by establishing higher basal levels of this AMP with which to counteract an incoming infection. This is not seen for the AMP *Drosomycin*, which again is consistent with our previous result that this AMP is not deployed against infection with *P. entomophila* (Fig. 4 and Suppl. Fig. 3).

With this we re-enforce the notion that the process of earlier AMP down-regulation could be an active response rather than a consequence of lower bacterial loads. This points to a potential role for disease tolerance mechanisms in this population through the prevention of AMP-induced immunopathology.

### A role for Disease Tolerance

Disease tolerance encompasses many distinct mechanisms of action, and describes multiple different physiological processes, which have in common the wide umbrella of maintaining host fitness without directly impacting pathogen load^6,77^. Maintenance of fitness in the face of stress-inducing or homeostatic conditions is assured by tissue damage control mechanisms^6,77^. Furthermore, these damage control responses can be activated in the event of an infection, either due to direct action of pathogens (i.e., injury-inducing toxins) or indirectly from immune-derived damages (i.e., immunopathology).

With this in mind, a putative role for disease tolerance was tested by measuring survival of the evolved populations to prolonged exposure to heat-killed bacteria. This protocol provided a means by which to test the response to damage, whether provoked by pathogenicity of the bacteria or by the host’s own response, independently of active pathogen elimination. Although we hypothesize that the heat-killing treatment was insufficient to denature all the virulence factors present in the suspension, we are yet to disentangle the relative contribution of virulence factors versus immunopathology to the longevity measurements. A similar longevity was observed for females and males of Control and BactOral after feeding on heat-killed bacteria, but an interesting trend appeared when comparing survival profiles (Fig. 6). We found that BactOral survived better to the stress of prolonged exposure to heat-killed *P. entomophila*, specifically under an evolutionarily relevant timeframe, that is, the period when flies reproduced during the selection experiment. Although both Control and BactOral individuals fully succumbed to the chronic exposure to heat-killed bacteria, significant differences in survival distinguish the two populations. Physiological responses measured approximately at 10 days post-infection (the age at which populations were selected to reproduce), can potentially represent evolutionary costs^78–81^, which are not the focus of this study.

Additionally, together with the gene expression data presented in Fig. 4, showing that BactOral represses the activation of immune effectors earlier upon infection, it is tempting to hypothesize that a similar process happened during the course of this experiment, and a role may be attributed to tighter control and negative regulation of immune over-activation.

The improved response of adapted flies to exposure to heat-killed bacterial substrate can be explained by a higher capacity to withstand damage, whether 1) provoked by their own immune response, or 2) caused by virulence factors present in inactivated bacterial supernatant. Additionally, process 1) could be correlated with the previously identified tighter immune repression. Disentangling the relative contribution of these mechanisms to the overall adapted phenotype would need further empirical characterization.

## CONCLUSION

In sum, we were able to identify resistance and disease tolerance mechanisms as targets of experimental evolution for increased survival to *P. entomophila* oral infection. Our observations align with previous work that identifies a positive genetic correlation between these two distinct immune defense strategies^82^. The increased survival of an experimentally adapted population to oral infection directly relates to its capacity to decrease bacterial loads after infection, independently of bacterial feeding rate, gut renewal capacity and bacterial defecation. Previous work found resistance to be targeted by selection upon immune challenge in *D. melanogaster*^79,83^. Our work adds to this approach by approaching the other mechanisms at play in immunity, namely disease tolerance and behaviour, in the framework of a single evolutionarily coherent system. We report one additional case of resistance evolution that exhaustively approaches the different processes by which immunity may evolve in populations fighting off persistent infections. In addition to resistance, we find a quantifiable role for disease tolerance in the evolved response of BactOral, revealing the complex nature of this adaptative process^82^, which is likely to rely also on improved tissue repair coupled with a more controlled deployment of damage-inducing immune effectors, such as AMPs^84,85^. This interplay between disease tolerance and resistance, underscores the complex mechanistic nature of immune response evolution with, undoubtedly, parallels in other host-pathogen systems. Future work will focus on identifying the specific mechanisms of disease tolerance and their physiological mode of action and interactions with resistance and their potential evolutionary repeatability.

## METHODS

### Maintenance of Drosophila populations under relaxed selection

Experimentally evolved Control and BactOral populations were initially derived from a highly outbred population of *D. melanogaster*, as described in previous studies ^19,46^. During experimental evolution, four replicates of each selection regime were maintained by orally infecting 310 females and 310 males per replicate, with a suspension of *P. entomophila* previously determined to cause approximately 66% of mortality in the starting population. After selection was terminated, populations were kept at high census (1500-2000 individuals per cage), allowing for maintenance of high genetic variability and optimal larval development. For all generations, flies were kept under constant temperature (25°C), humidity (55–65%) and light-darkness cycle (12:12), and fed with a standard cornmeal-agar medium, consisting of 4.5% molasses, 7.5% sugar, 7% corn-flower, 2% granulated yeast extract, 1% agar and 0.25% nipagin, mixed in distilled water. Control and BactOral were under selection for 24 generations and under relaxed selection for approximately 80 generations, after which one replicate of each regime was singled out to perform mechanistic experiments.

Each generation cycle lasted approximately 3 weeks, during which flies eclosed and remained undisturbed until they were 7-8 days old. At this time, fresh food was placed in the cages for 2 days, which allowed for oviposition of retained eggs. After this period (when flies were 9-11 days old), similarly to the protocol applied for selection, new fresh food was placed in the cages for controlled egg-lays (1-3 hours long) and establishment of the next generation.

### Pathogen culture and infections

Oral infection protocol with *P. entomophila* (Rifampicin resistant strain kindly provided by Bruno Lemaitre) was adapted from the one used in Martins and Faria et al^19^. Briefly, single bacterial colonies were grown in kick-start cultures (5mL of LB) for approximately 8 hours after which they were transferred into larger volumes (1 L of LB) for overnight growth, both periods at 29°C. After centrifugation and resuspension of bacterial pellets, concentration was adjusted to OD600= 100 and finally diluted 1:1 with a 5% sucrose solution. For infection, 3-5 days-old flies were separated by sex into groups of twenty with the use of CO2 (at least 24 hours before the start of the experiments) and allowed to feed on a filter disk embedded with the bacterial solution described above for 24 hours (for survival assays) or for the amount of time specific to each experiment (see Results). After this period, infected flies were flipped onto clean food and survival was scored daily. Controls were fed a 1:1 LB/PBS:5% sucrose solution.

For the priming experiment, a solution of *P. entomophila* at OD600= 100 was previously heat-killed for 1 hour at 55°C, mixed 1:1 with a 5% sucrose solution and fed to flies for 2 hours, in a similar way as described above for regular oral infection. Efficiency was confirmed by plating the heat-killed culture and confirm absence of bacterial growth. After this time, flies were transferred into tubes containing a live solution of *P. entomophila* for 24 hours, after which they were changed back to clean food and their survival scored.

For the experiment with prolonged exposure to heat-killed *P. entomophila* a similar bacterial preparation and heat-killing protocol were performed as described above, but the final dead bacteria suspension was mixed 1:1 with standard fly food and dispensed into fly vials. To the final solution, extra agar powder was added to account for viscosity and to optimize texture and left to cool down and solidify for at least 2 hours, before being fed to flies. The same protocol was performed for the control treatment, in which fly food was mixed with sterile 1X PBS. Exposure to these treatments was constant throughout the experiment, during which replicates of 15 females and 15 males were co-housed in vials, exclusively feeding on the dead-bacteria or PBS mixtures. Survival scoring began at the start of exposure and lasted 14 days. During the treatment, flies were frequently flipped onto fresh food mixture and survival scoring continued daily. Overall efficiency of the bacterial heat-killing protocol was tested by culturing the heat-killed suspension and confirm the absence of bacterial growth, prior to storage at -20 °C, which anteceded the mixing with the food or sucrose solution.

Survival analysis was done for the multiple experiments by fitting data with Cox Proportional Hazards models (with *coxme* or *coxph* functions from the *survival* package^86^) using population, treatment, and sex as fixed factors and replicate (or replicate nested within block) as a random factor. To assess the effect sizes of each factor, type 3 ANOVAs (*Anova* from the *car* package^87^) were ran on the survival models. Additionally, multiple comparisons with *tukey* adjustments were done on all the models to disentangle which specific pairwise comparisons were significantly different (*emmeans* function^88^). Finally, the *cld* function of the *multcomp* package^89^ (with *sidak* confidence-level adjustments) was used to summarize the results of the multiple comparisons across conditions. Survival models used were:

- immunocompetence (Fig. 1)- *coxme*(∼Population*Treatment*Sex+(1|Replicate));
- priming (Fig. 5)- *coxme*(∼Population*Treatment*Sex+(1|block/Replicate)) and
- immune over-activation (Fig. 6)- *coxme*(∼Population*Treatment*Sex+(1|Replicate), data = survival_HKexposure).

### Bacterial defecation assay

To have a measure of the rate of gut purge/defecation a change was introduced to the bacterial infection protocol, whereby the feeding period was reduced to a 3-hour period. After this time, flies were surface sterilized and separated individually into plastic spectrophotometry cuvettes, previously filled with fly food. After 6 hours (6 hours.), the same flies were changed into fresh cuvettes where they remained for an additional 12 hours (18 hours.), after which they were discarded. Bacterial quantification was performed for both timepoints by washing the interior surface of the cuvettes with PBS and plating, as described below.

Statistical analysis on number of defecated bacteria was done by fitting a zero-inflated on the CFU counts and zeros (using the *zeroinfl* function^90^ from the *pscl* package^91^). This model allowed for the fitting of a negative-binomial distribution on the counts data and of a binomial distribution on the zeros (*zeroinfl* (formula= Counts ∼ Population * Sex * Timepoint | Population * Timepoint, dist= “negbin”)). To assess differences between specific pairwise comparisons we used *emmeans* with *tukey* adjustments, on both distributions of the zero-inflated model. In the instances where flies died during either timepoint, their count at that time was removed from further analysis and the linear regression was not included in Figure 2B.

### Quantification of food intake by CApillary FEeder Assay - CAFE

To measure the amount of bacterial solution ingested by single flies, an adapted CAFE^47^ assay was performed. We substituted the food of small fly vials with agarose, to ensure humidity, and placed a glass capillary in each lid, filled with 5% sucrose solution. We allowed individual flies to habituate to this experimental setup for 24 hours before the infection. After this period, capillaries filled with *P. entomophila* suspension (prepared as described before) were given to individual females to feed on and consumption estimated, at different timepoints. To facilitate scoring in capillaries, the bacterial solution was mixed with blue food colouring.

Modelling of cumulative amounts of bacteria eaten over time was done with a Generalized Linear Model with a Gamma distribution, (*glm*(Quantity ∼ Population*Timepoint)) using the *glm* function from the base *stat* R package^93^. For better visualization of feeding over time, linear regressions were drawn between mean values of microliters eaten cumulatively by each population at each timepoint, using the *stat_poly_eq* function of the *ggpmisc* package^94^. Finally, multiple comparisons were done with *emmeans*^88^, using *tukey* adjustments, to disentangle differences between populations across timepoints.

### Measurement of gut length

Females were individually dissected in PBS 1X, and their guts removed for imaging by adapting a protocol described in^96^. Dissections started by fixing flies in the head with a Minutien pin and gradually separating the gut from the remaining tissues, in a posterior-to-anterior direction. After the gut was exposed, we added a few drops of a solution of 5% Acetic Acid and 50%EtOH, which slightly whitens the tissues, allowing for better visualization and a slight tissue fixation. After having isolated the guts, they were individually transferred to a coverslip and photographed using a uEye camera installed on an Olympus SZX7 stereoscope.

Images were processed in FIJI v2.0.0 and measurements were collected by drawing consecutive transects along the total length of guts, as illustrated in Supplementary figure 2A. Each individual gut measurement was repeated 3 times, to account for errors in the manual drawing of the transects and averaged to obtain a final gut length in millimeters (mm).

Differences in lengths of guts between populations were analysed with a Linear Mixed-Effects Model (*lmer*(Length ∼ Population * Treatment + (1|Block)) with *lmer* from the *lmerTest* package^97^. The model was followed by multiple comparisons with *emmeans*^88^, with a *tukey* adjustment, for assessment of individual pairwise differences.

### Quantification of bacterial loads

To quantify bacterial loads, we adapted an established protocol^19,92^. In short, individual flies were surface sterilized (washed in EtOH 70%, bleach 60% and two washes with MiliQ water) and placed in 96 deep-well plates, where they were homogenized with glass beads in 50 µL of LB using a TissueLyser II (Qiagen). Individual samples were serially diluted 5 times with 1:10 dilution steps, plated as 4 µL droplets in LB+Rifampicin, and left to grow overnight at 29°C to optimize colony size for counting. For the bacterial defecation assay, cuvettes that contained flies for each of the timepoints were filled with 1mL of PBS and incubated at room temperature for 15 minutes, after which each sample was diluted up to 4 times (1:5 dilutions) and platted as droplets in LB+Rifampicin.

Analysis of the time-course of infection progression was performed by dividing the data obtained from the bacterial quantification into “counts” and “zeros” and using different models to examine their dynamics in our evolved populations. For the counts, the modelling of the mean of the load dynamics during the time-course was done using a sigmoidal function that considers four parameters: *a* the log (initial load), *k* the timepoint at which the load is half of the maximum value (in this case, *a*), *h* the slope at the *k* timepoint (steepest slope of the curve), and *b* the log (final load). A linear model was applied to each of the parameters which was fitted using a Monte Carlo Markov Chain approach. To estimate the impact of Sex and Population on the likely distribution of the parameters, a normal error structure was assumed, using Control females as reference. For the “zeros”, a modified logistic model with four parameters was ran: *a0* initial fraction of zeros, *k0* rate of zero disappearance, *k1* rate of zero re-appearance, *t0* timepoint at which zeros re-appear and *a1* final fraction of zeros. Similarly, to the “counts” analysis, linear models were used to assess the impact of Sex and Population on all four parameters.

### RNA extractions and quantitative PCR

For RNA extractions, flies from Control and BactOral population were infected as described above and, at different timepoints, pools of 10 females were collected and homogenized in 500 µL of Trizol. RNA extractions were performed using a Direct-zol™ RNA MiniPrep w/ Zymo-Spin™ IIC Columns. After precipitation, a DNase I (RQ1 RNASE-FREE DNASE 1* from Promega®) treatment was done to all samples, followed by reverse transcription using the Thermo Scientific® RevertAid H Minus cDNA kit. cDNA was finally diluted 1:5 for qPCR.

For quantification of gene expression, qPCRs were performed using SYBRTM Green Master Mix (Thermo Scientific®) and reactions ran on 384-well plates (Applied Biosystems®). The PCR conditions used in all experiments were: initial denaturation/ enzyme activation, 95°C for 10’; followed by 45 cycles of denaturation, 95°C for 10”; annealing, 60°C for 10”; extension, 72°C for 30”. Sequences of primers used for qPCRs are shown in Supp. Table. 1. Either EIF2 (eIF-2alpha) or Rpl32 were used as reference.

Analysis of gene expression differences was done with the relative quantification technique (DDCt)^95^. Briefly, to the average of technical replicates for each candidate gene we subtracted the average of the Ct values of the respective sample’s house-keeping gene (EIF2 or Rpl32) (DCt) and normalized this value to the DCt of the respective reference condition. For the time-course analysis, the reference was the “Unchallenged” timepoint while for comparisons between “Unchallenged” conditions among populations, Control was used as reference. Finally, we used reverse logarithm to transform the final gene expression values into fold change levels (2^-DDCt^) and analysed the Log2(foldchange) differences across Populations and/or Timepoints.

Statistical analysis on gene expression was done by running Linear Models (from base R stats^93^) on the Log2(foldchange) values of each individual gene (*lm*(logfold_AMP ∼ Population * Timepoint), followed by an ANOVA (type three) from the car package^87^. To individualize significant pairwise comparisons, *emmeans* was ran on the linear models (with a Benjamini-Hochberg adjustment). Finally, the *cld* function of the multcomp package^89^ (with sidak confidence-level adjustments) was used to summarize the results of the multiple comparisons across different conditions.

### Statistical analysis and graphical representations

All datasets (excluding survival data) were tested for Normality before further analysis, using the Shapiro-Wilk Test (*shapiro.test* function of the *stat* R package), for determination of the most suitable type of statistical test to perform in each case. All model formulas used for statistical analysis are discriminated throughout the Results sections. All graphical representations were performed using the *ggplot2* package^98^.

Statistical analysis and graphics for all experiments, excluding analysis of the bacterial loads time-course modelling analysis (Fig. 5), were done on R version 4.2.1, through RStudio^99^ version 1.3.959. For the bacterial dynamics time-course, models were implemented and fitted in Python^100^, using the probabilistic programming language package *pymc3*^101^ version 3.11.

Sensitivity (power) calculations for the CAFE and qPCR datasets was calculated using G*Power 3.1 (https://www.psychologie.hhu.de/arbeitsgruppen/allgemeine-psychologie-und-arbeitspsychologie/gpower).

Sensitivity analysis for gut length dataset was performed by simulation, using Python packages *scipy*, *numpy*, and *statsmodels*. Briefly: a synthetic dataset of the same sample sizes and structure as the real data but with a specified effect in both factors (Treatment and Genotype) was generated. Multiple linear regression was then performed and the significance (*p-value*) of both factors was recorded. This was performed 10^5^ times, and the fraction of significant tests was recorded (power, independently for both factors). A numerical search for the minimum effect sizes that achieve 80% power on both factors was then performed.

## ACKNOWLEDGEMENTS

We thank Dr Bruno Lemaitre for stocks, Liliana Vieira and Sandra Crisóstomo for technical assistance, André Barros and David Duneau for help with analyses, and Elvira Lafuente, Diogo Roque and Patrícia Duarte for critical comments and discussions that improved the manuscript. This work was supported by Instituto Gulbenkian de Ciência/Fundação Calouste Gulbenkian and FCT-Fundação para a Ciência e a Tecnologia (Portugal) UID/Multi/04555/2013 and UIDB/00329/2020. TFP (BD/128432/2017 and COVID/BD/151645/2021) and to PAA (BD/06404/2020) were supported FCT—Fundação para a Ciência e a Tecnologia (Portugal); and by C.S. CONGENTO, project LISBOA-01-0145-FEDER-022170, co-financed by Lisboa Regional Operational Programme (Lisboa 2020), under the Portugal 2020 Partnership Agreement, through the European Regional Development Fund (ERDF), and Foundation for Science and Technology (Portugal).

## DECLARATION

The authors declare that this work is not under review elsewhere and that none of its authors has any conflict of interest to declare.

## DATA AVAILABILITY

All relevant data are within the manuscript and its Supporting Information files.

## AUTHOR CONTRIBUTIONS

Conceptualization: Tânia F. Paulo and Élio Sucena.

Data curation: Tânia F. Paulo

Formal analysis: Tânia F. Paulo, Tiago Paixão

Funding acquisition: Élio Sucena.

Investigation: Tânia F. Paulo, Priscilla A. Akyaw and Élio Sucena.

Methodology: Tânia F. Paulo, Priscilla A. Akyaw, Tiago Paixão and Élio Sucena.

Resources: Élio Sucena and Tiago Paixão.

Supervision: Élio Sucena.

Visualization: Tânia F. Paulo.

Writing – original draft: Tânia F. Paulo and Élio Sucena.

Writing – review & editing: Tânia F. Paulo, Priscilla A. Akyaw, Tiago Paixão and Élio Sucena.

## SUPPLEMENTARY FIGURE

**Supplementary Figure 1.**
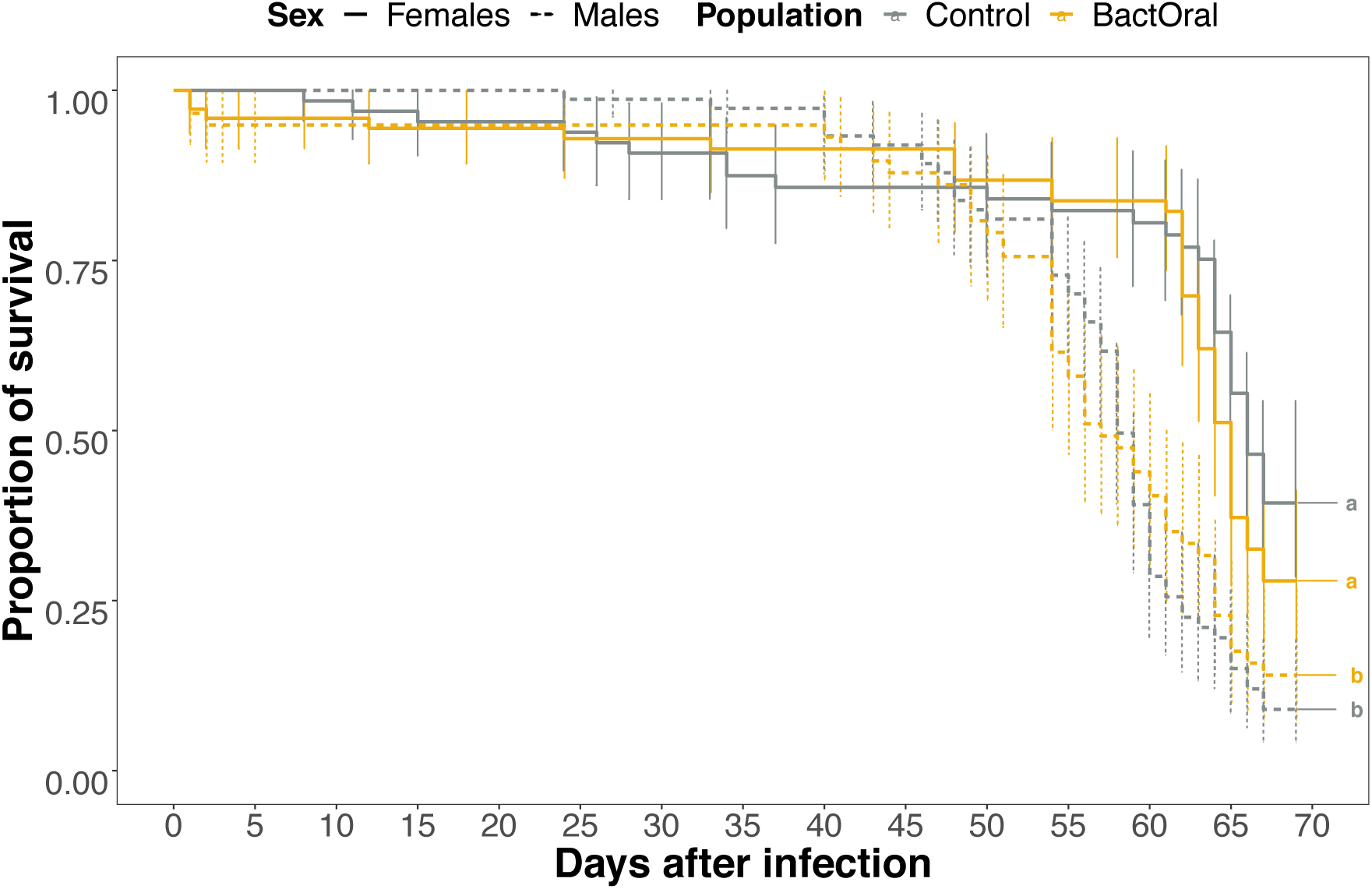
Longevity of BactOral and Control populations. Females (full lines) and males (dashed lines) of Control (grey) and BactOral (yellow) populations, are shown. No differences can be found between populations within sex (Uninfected Control females versus Uninfected BactOral females: z.ratio= 1.214, p= 0.6180; Uninfected Control males versus Uninfected BactOral males: z.ratio= 0.615, p= 0.9272). Results from multiple comparisons are shown as lower-case letters in the plots.

**Supplementary Figure 2.**
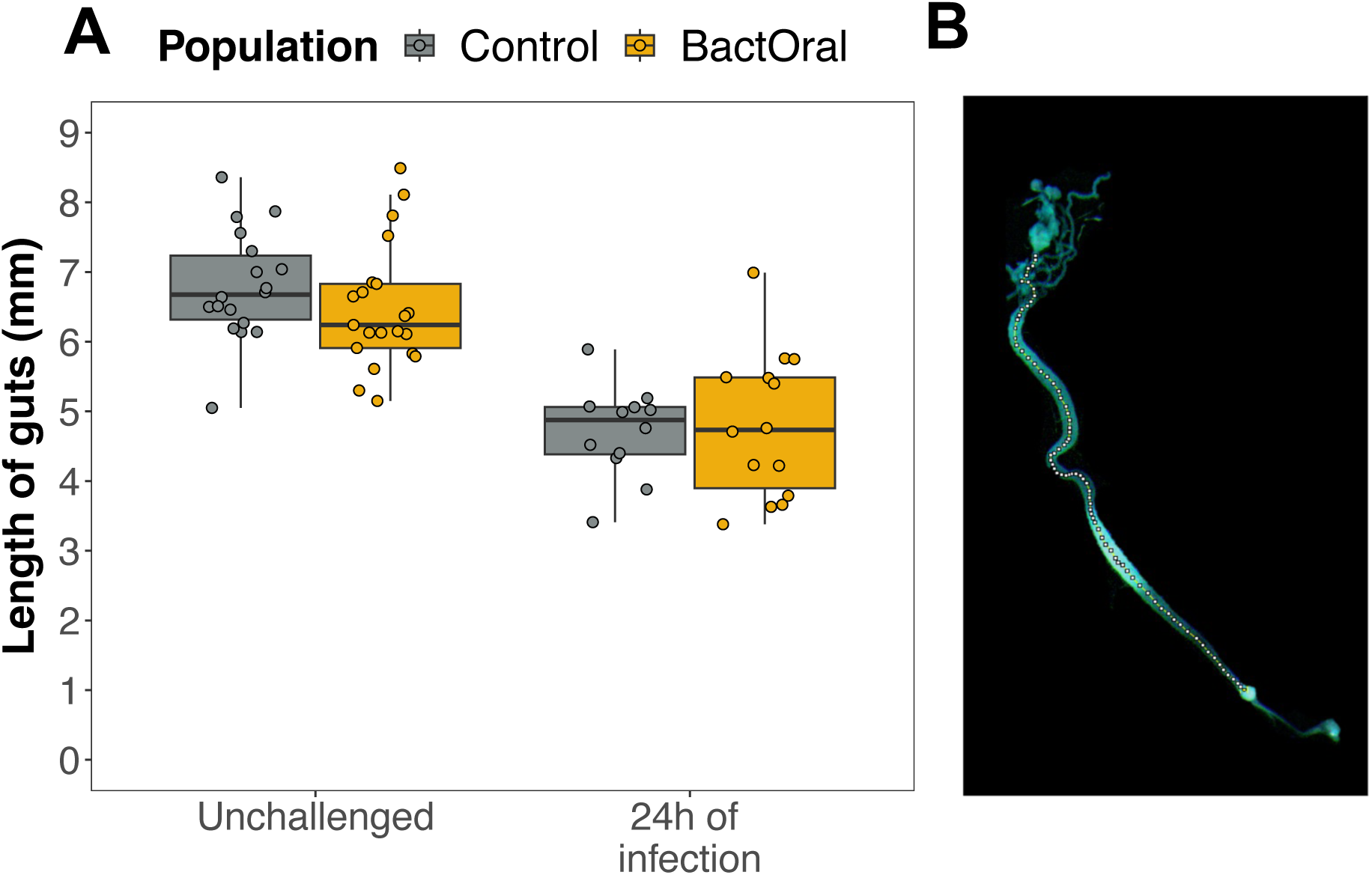
No detectable differences in gut structure between Control and BactOral after oral infection with *P. entomophila*. Females of Control (grey) and BactOral (yellow) were fed normal food (“Unchallenged”) or orally infected with *P. entomophila* for 24 hours, after which their **A)** guts were dissected and photographed **B)**. Gut length (in mm) was measured by drawing consecutive longitudinal transects along the guts thrice and averaging those measurements to reach final length per sample. **A)** Exposure to bacteria shortens the guts significantly in both populations (Control before infection *versus* Control 24 hours post-infection: *t*= 6.274, *p*<0.0001; BactOral before infection *versus* BactOral 24 hours post-infection: *t*= 5.453, *p*<0.0001). No differences were detected in gut lengths between populations within each timepoint (Without infection: BactOral *versus* Control – *p*= 0.6719; after 24 hours of infection – *p*= 0.9925), although a significant effect of treatment was observed in both Control (Without infection *versus* after 24 hours of infection – *p*< 0.0001) and BactOral (*p*< 0.0001), showing that gut shortening due to infection affected both regimes in a similar manner. **B)** Dissected guts of females from Control and BactOral populations were photographed and scaled for posterior length quantification. Manual transects were drawn longitudinally along the guts (as shown above) and total size of the cumulative transects was calculated in FIJI. Three independent measurements were done on each gut and averaged to obtain a final value per sample (Supp. Fig. 2 A)). Samples sizes are shown at the top of the panel.

**Supplementary Figure 3.**
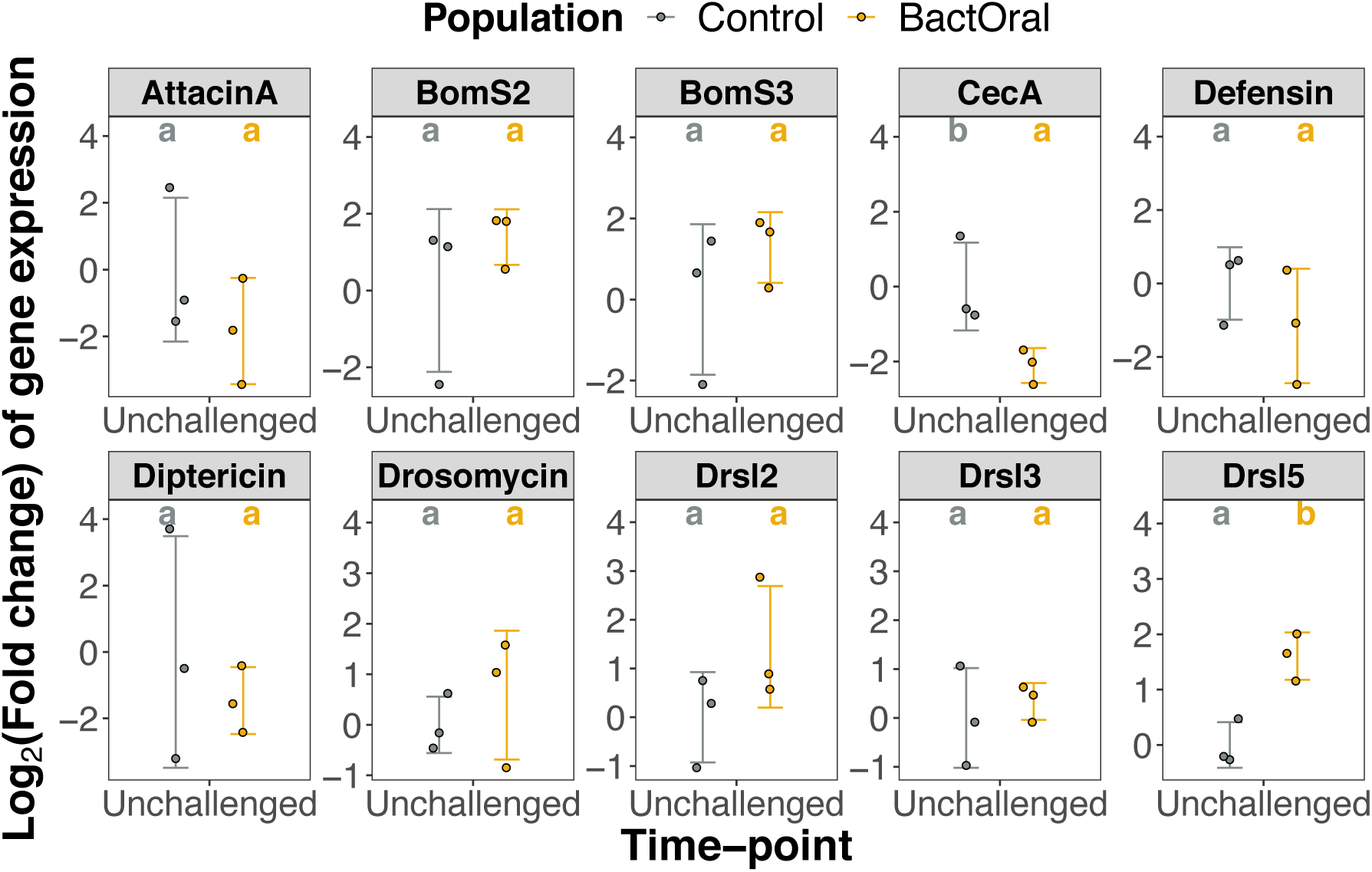
AMP expression differences between populations under homeostatic conditions. Guts from Control (grey) and BactOral (yellow) female flies (3 samples containing 10 guts, each) were dissected at different timepoints after oral infection with *P. entomophila* and relative quantification of gene expression was performed. Expression levels of each gene in BactOral samples were normalized to expression levels of Control samples. Log2(FoldChange) of these differences were fitted on a linear model with Population as factor and followed by multiple comparisons (*emmeans*) to uncover pairwise differences. Results from these comparisons are shown in the letters above the plots. There are significantly lower levels of CecropinA (CecA) (*t*= 2.892, *p*= 0.0445) and significantly higher levels of *Drosomycin-like 5* (Drsl5) (*t*= -4.686, *p*= 0.0094) being expressed in guts of BactOral flies under unchallenged conditions.

**Supplementary Figure 4.**
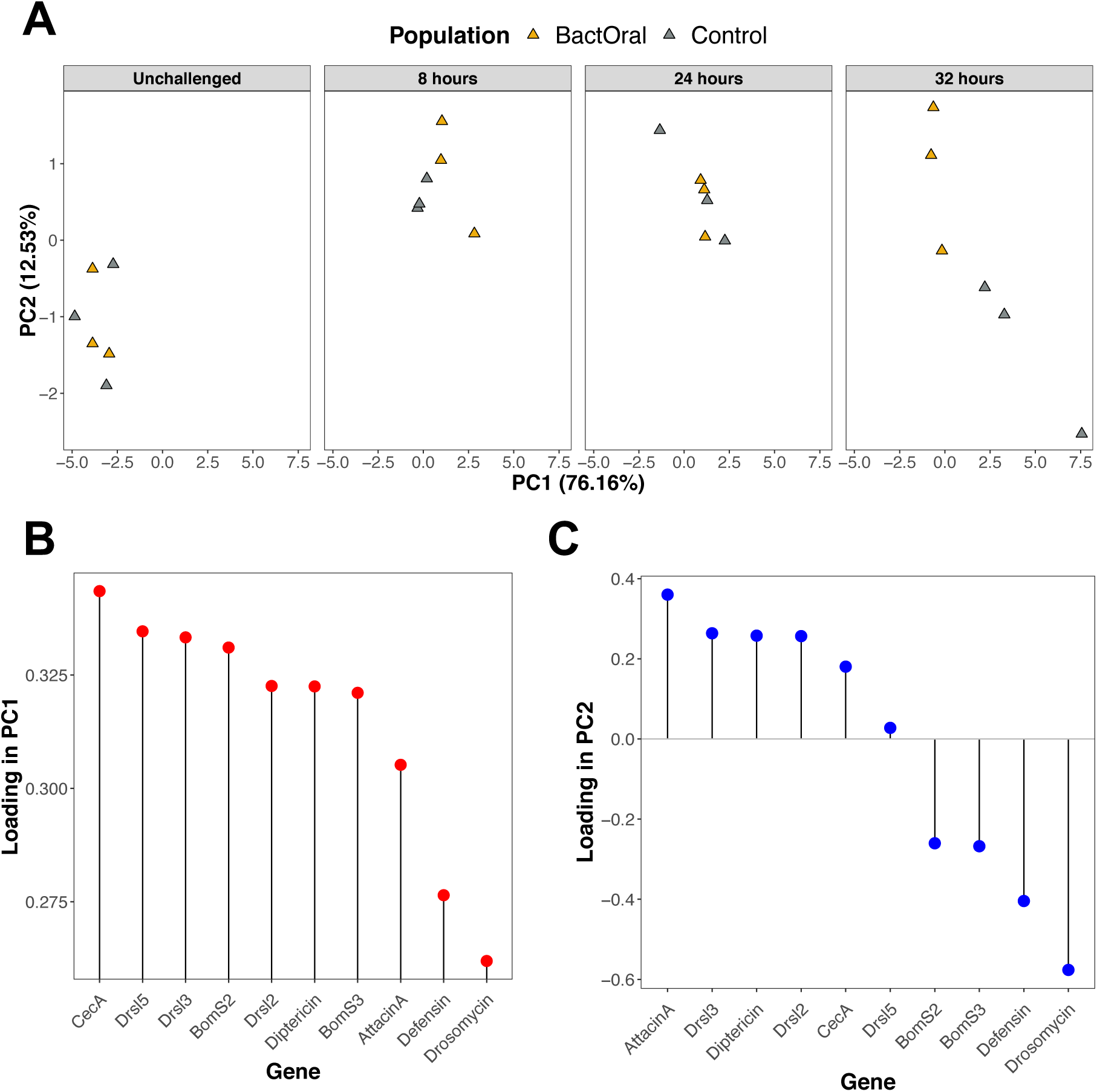
Principal Component Analysis on relative gene expression levels in evolved populations. (A) Relative AMP expression levels of Control and BactOral flies upon oral infection with *P. entomophila* showed significant differences at various timepoints (see Fig. 4 and Supp. Fig. 2). **(B-C)** Relative influence of each AMP to the establishment of PC1 and PC2 (see Supp. Fig. 3 A). Importantly, Cecropin A1 represents the largest contributor to PC1 (**B**) whereas Drosomycin represents the largest contributor, although negatively correlated, to the variation in PC2 (**C**).

**Supplementary Figure 5.**
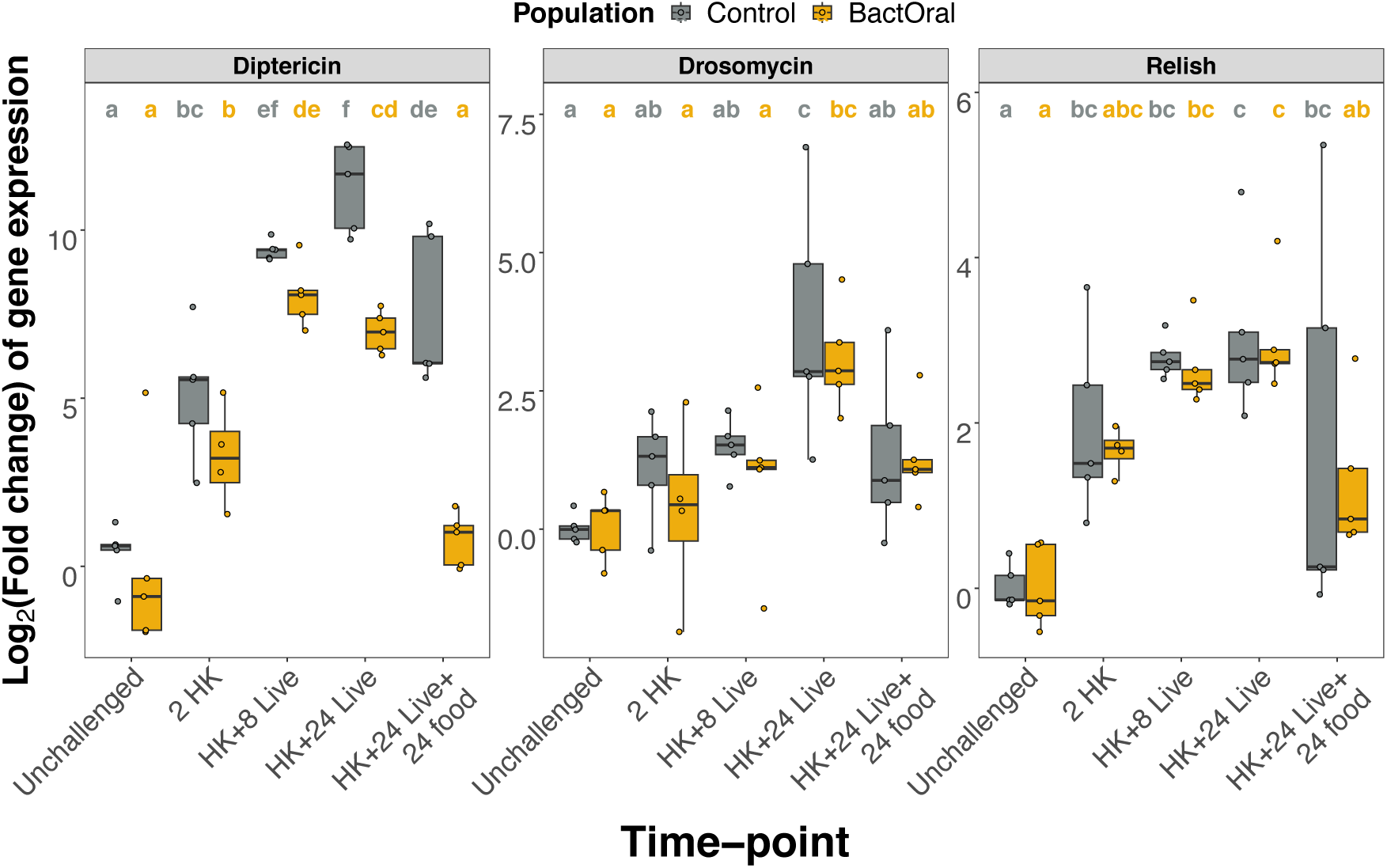
**Quantification of immunity-related gene expression upon artificial immune stimulation and infection**. RNA from whole-females from Control (grey) and BactOral (yellow) (4 samples containing 10 flies, each) was extracted at different timepoints. Relative gene expression, normalized to housekeeping gene *rpl32,* was performed for *Diptericin*, *Drosomycin* and *Relish*. Results from these comparisons are shown in the letters above the plots. To test for individual pairwise differences, relative Log2 (FoldChange) expression levels of each gene were fitted onto a linear model with Population and Timepoints as factors and followed by multiple comparisons (*emmeans*). No differences are observed between non-challenged individuals or between individuals after 2 hours of exposure. However, for Diptericin Unchallenged versus 2 HK show differences in both populations (Control: *t*= -4.778, *p*= 0.0001; BactOral: *t*= -3.122, *p*= 0.0052).

## SUPPLEMENTARY TABLES

**Supplementary Table 1.**
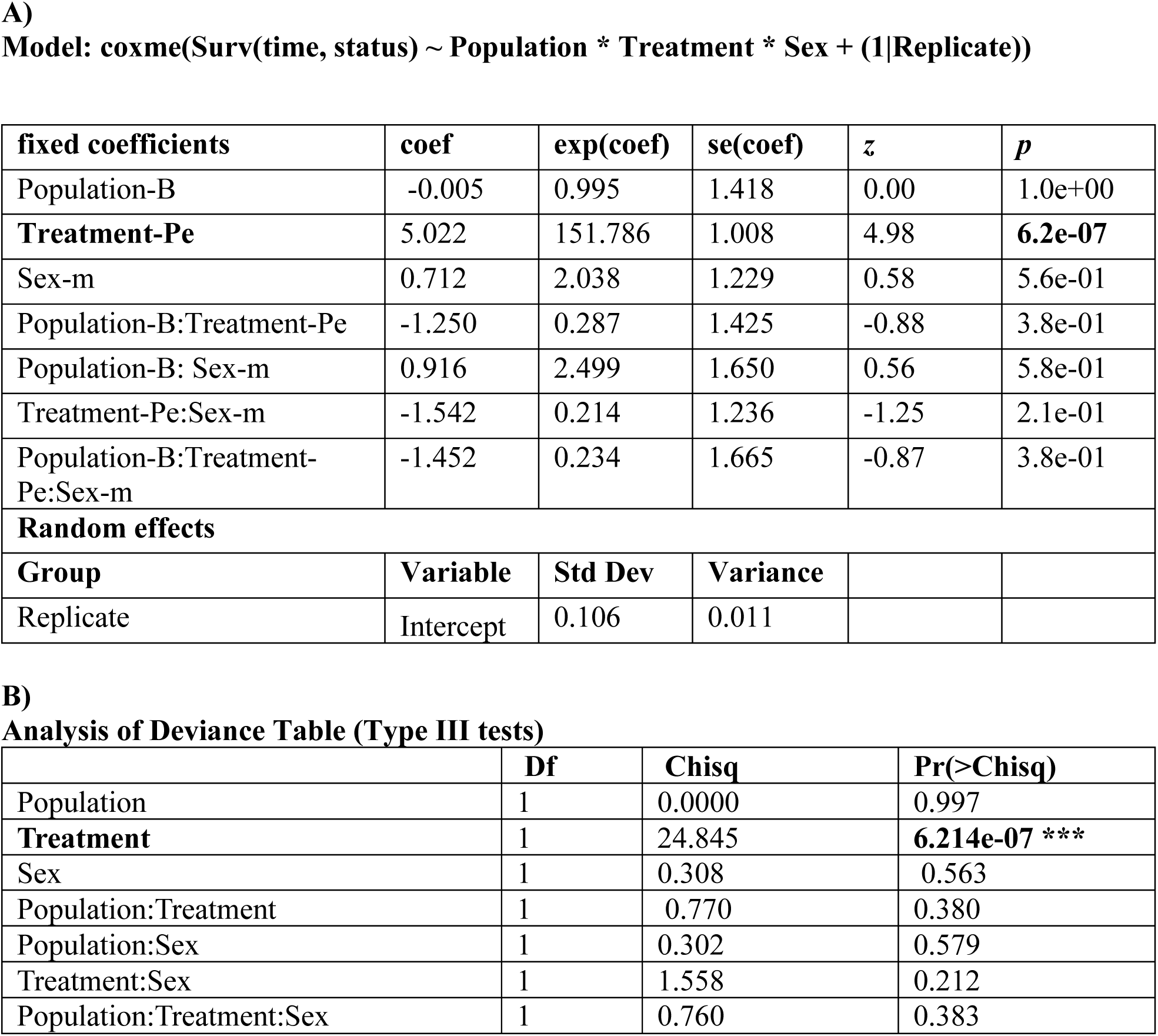
Statistical analysis of survival after oral infection with *P. entomophila* (immunocompetence, shown in Fig. 1). A) Model estimates and B) ANOVA estimates for survival analysis testing immunocompetence of Control and BactOral populations.

| fixed coefficients | coef | exp(coef) | se(coef) | z | p |
| --- | --- | --- | --- | --- | --- |
| Population-B | -0.005 | 0.995 | 1.418 | 0.00 | 1.0e+00 |
| **Treatment-Pe** | 5.022 | 151.786 | 1.008 | 4.98 | **6.2e-07** |
| Sex-m | 0.712 | 2.038 | 1.229 | 0.58 | 5.6e-01 |
| Population-B:Treatment-Pe | -1.250 | 0.287 | 1.425 | -0.88 | 3.8e-01 |
| Population-B: Sex-m | 0.916 | 2.499 | 1.650 | 0.56 | 5.8e-01 |
| Treatment-Pe:Sex-m | -1.542 | 0.214 | 1.236 | -1.25 | 2.1e-01 |
| Population-B:Treatment-Pe:Sex-m | -1.452 | 0.234 | 1.665 | -0.87 | 3.8e-01 |
| **Random effects** |  |  |  |  |  |
| Group | Variable | Std Dev | Variance |  |  |
| Replicate | Intercept | 0.106 | 0.011 |  |  |

**Supplementary Table 1. Statistical analysis of survival after oral infection with *P. entomophila* (immunocompetence, shown in Fig. 1).**
|  | Df | Chisq | Pr(>Chisq) |
| --- | --- | --- | --- |
| Population | 1 | 0.0000 | 0.997 |
| **Treatment** | 1 | 24.845 | **6.214e-07 \*\*\*** |
| Sex | 1 | 0.308 | 0.563 |
| Population:Treatment | 1 | 0.770 | 0.380 |
| Population:Sex | 1 | 0.302 | 0.579 |
| Treatment:Sex | 1 | 1.558 | 0.212 |
| Population:Treatment:Sex | 1 | 0.760 | 0.383 |

**Supplementary Table 2.** Statistical analysis of survival under uninfected conditions (Longevity, shown in Supp. Fig. 1). A) Model estimates and B) ANOVA estimates for survival analysis testing longevity of Control and BactOral populations.

|  | **coef** | **exp(coef)** | **se(coef)** | **z** | **Pr(>\|z\|)** |
| --- | --- | --- | --- | --- | --- |
| Population-B | 0.271 | 1.312 | 0.224 | 1.214 | 0.225 |
| **Sex-m** | 1.069 | 2.912 | 0.215 | 4.970 | **6.7e-07 \*\*\*** |
| Population-B: Sex-m | -0.389 | 0.677 | 0.295 | -1.321 | 0.186 |

**Supplementary Table 2. Statistical analysis of survival under uninfected conditions (Longevity, shown in Supp. Fig. 1).**
|  | **Chisq** | **Df** | **Pr(>Chisq)** |
| --- | --- | --- | --- |
| Population | 1.489 | 1 | 0.222 |
| **Sex** | 26.067 | 1 | **3.298e-07 \*\*\*** |
| Population:Sex | 1.759 | 1 | 0.185 |

**Supplementary Table 3.**
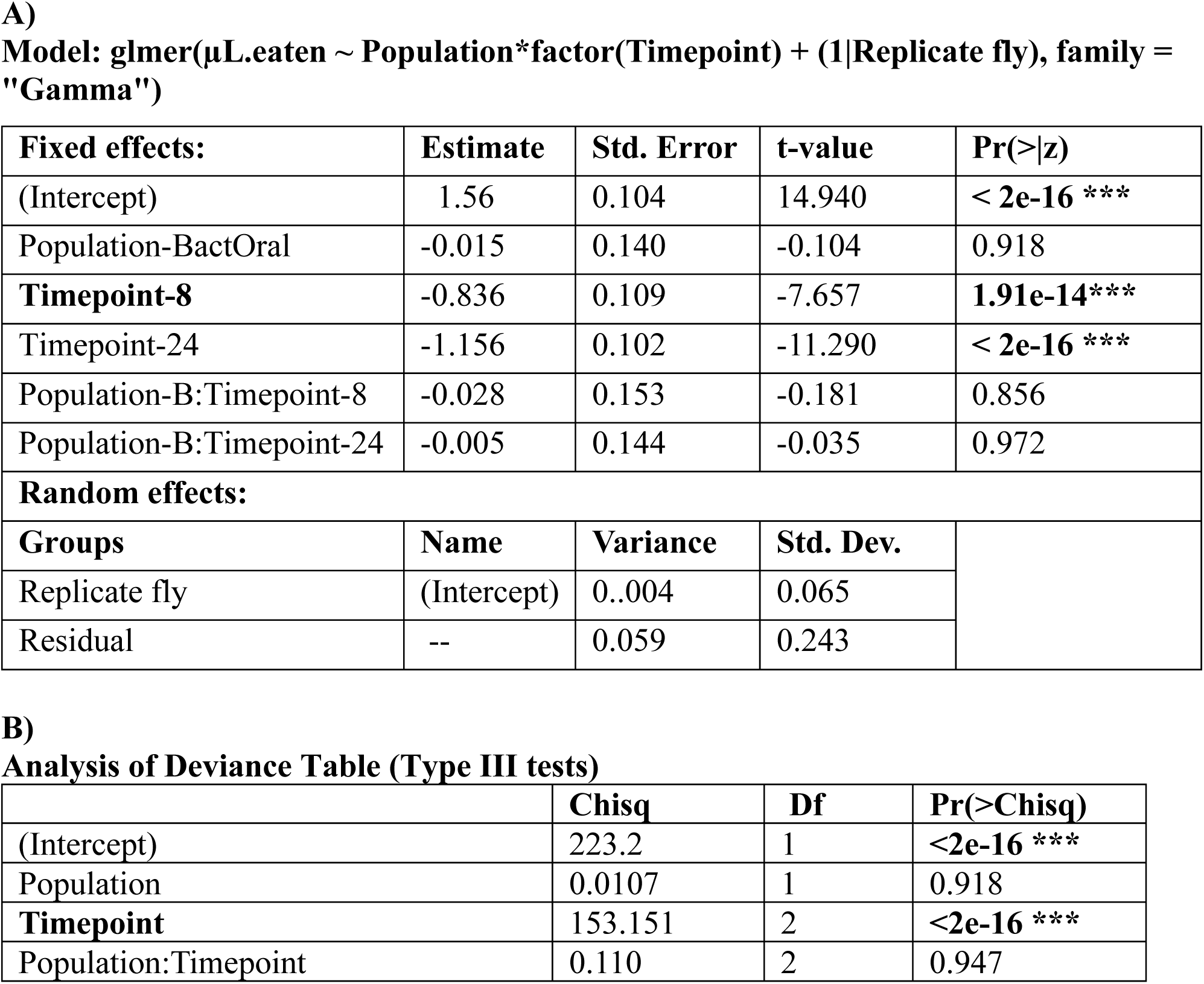
Analysis of the cumulative amounts of bacteria ingested by evolved populations (CAFE assay, shown in Fig. 2 A). A) Model estimates and B) ANOVA estimates for the generalized linear model testing the effects of population and timepoint on ingested bacterial mixture.

| Fixed effects: | Estimate | Std. Error | t-value | Pr(>\|z) |
| --- | --- | --- | --- | --- |
| (Intercept) | 1.56 | 0.104 | 14.940 | < 2e-16 *** |
| Population-BactOral | -0.015 | 0.140 | -0.104 | 0.918 |
| **Timepoint-8** | -0.836 | 0.109 | -7.657 | **1.91e-14\*\*\*** |
| Timepoint-24 | -1.156 | 0.102 | -11.290 | < 2e-16 *** |
| Population-B:Timepoint-8 | -0.028 | 0.153 | -0.181 | 0.856 |
| Population-B:Timepoint-24 | -0.005 | 0.144 | -0.035 | 0.972 |
| Random effects: |  |  |  |  |
| Groups | Name | Variance | Std. Dev. |  |
| Replicate fly | (Intercept) | 0.004 | 0.065 |  |
| Residual | -- | 0.059 | 0.243 |  |

Supplementary Table 3. Analysis of the cumulative amounts of bacteria ingested by evolved populations (CAFE assay, shown in Fig. 2 A).
|  | Chisq | Df | Pr(>Chisq) |
| --- | --- | --- | --- |
| (Intercept) | 223.2 | 1 | <2e-16 *** |
| Population | 0.0107 | 1 | 0.918 |
| **Timepoint** | 153.151 | 2 | **<2e-16 \*\*\*** |
| Population:Timepoint | 0.110 | 2 | 0.947 |

**Supplementary Table 4.** Analysis of quantities of defecated bacteria by evolved populations (CFUs, shown in Fig. 2 B)). A) Model estimates and for the zero-inflated model testing the effects of population, sex and timepoint on amount of defecated live bacteria and instances of non- measurable bacteria (zeros).

| Count model coefficients (negbin with log link) |  |  |  |  |
| --- | --- | --- | --- | --- |
|  | Estimate | Std. Error | z value | Pr(>\|z\|) |
| **(Intercept)** | 13.721 | 0.482 | 28.447 | **<2e-16 \*\*\*** |
| Population-C | 0.816 | 0.616 | 1.324 | 0.185 |
| Sex-m | 0.047 | 0.647 | 0.073 | 0.942 |
| Timepoint-18 | -0.251 | 0.699 | -0.359 | 0.720 |
| Population-C:Sex-m | -0.910 | 0.840 | -1.083 | 0.279 |
| Population-C:Timepoint-18 | -0.579 | 0.905 | -0.640 | 0.522 |
| Sex-m:Timepoint-18 | -0.950 | 0.911 | -1.043 | 0.297 |
| Population-C:Sex-m:Timepoint-18 | 2.171 | 1.204 | 1.803 | 0.071 |
| **Log(theta)** | -1.031 | 0.106 | -9.700 | **<2e-16 \*\*\*** |
| Zero-inflation model coefficients (binomial with logit link): |  |  |  |  |
|  | Estimate | Std. Error | z value | Pr(>\|z\|) |
| **(Intercept)** | -0.828 | 0.351 | -2.357 | **0.018 \*** |
| Population-C | -3.004 | 1.248 | -2.407 | **0.016\*** |
| Timepoint-18 | -18.771 | 3347.432 | -0.006 | 0.996 |
| Population-C:Timepoint-18 | 18.876 | 3347.433 | 0.006 | 0.996 |

**Supplementary Table 5.**
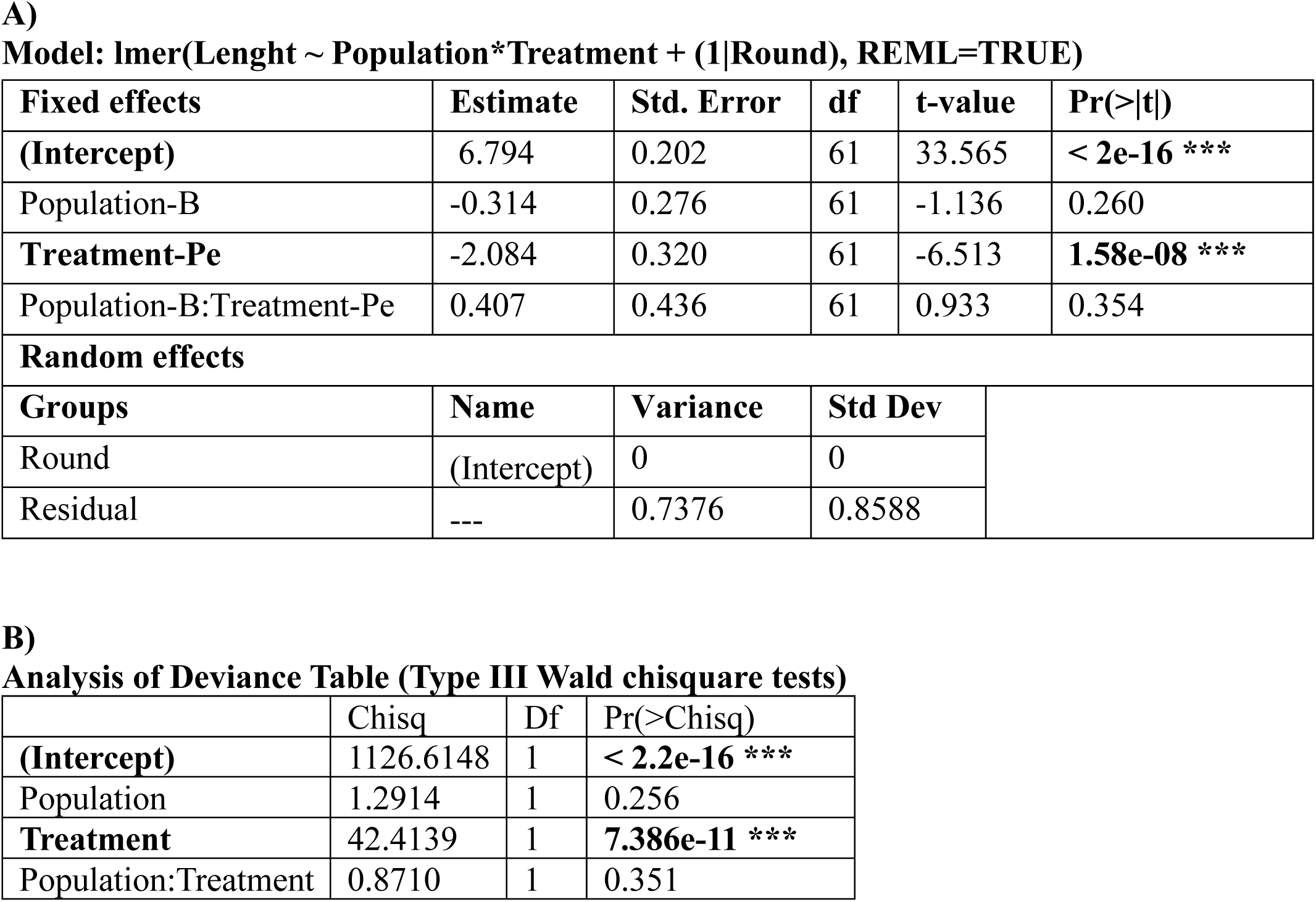
Analysis of total gut lengths of females of evolved populations before and after 24 hours of infection (shown in Supp. fig. 1 A)). A) Model estimates and B|) ANOVA estimates for the linear model testing the effects of population and treatment on total length of guts.

**Supplementary Table 6.**
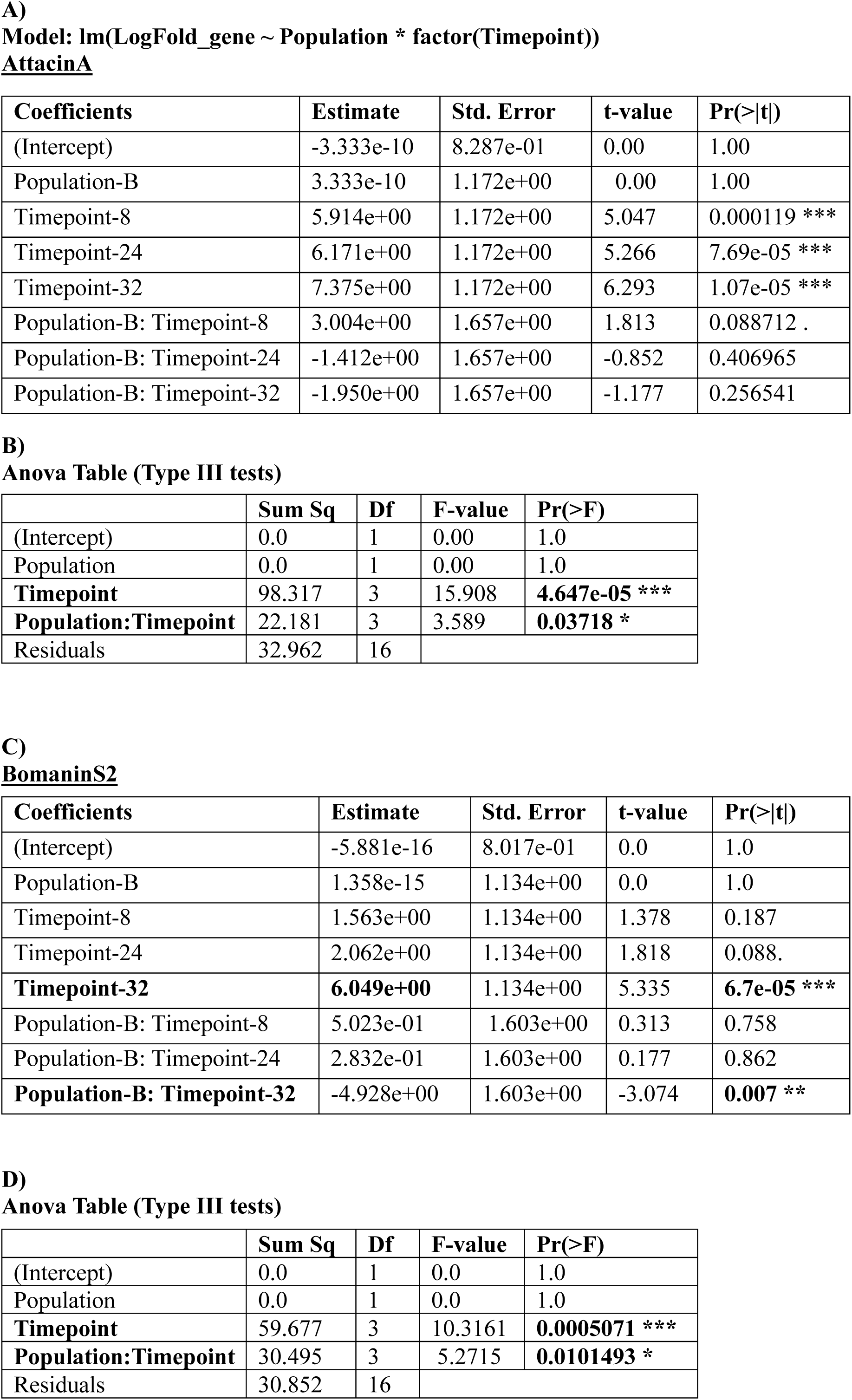
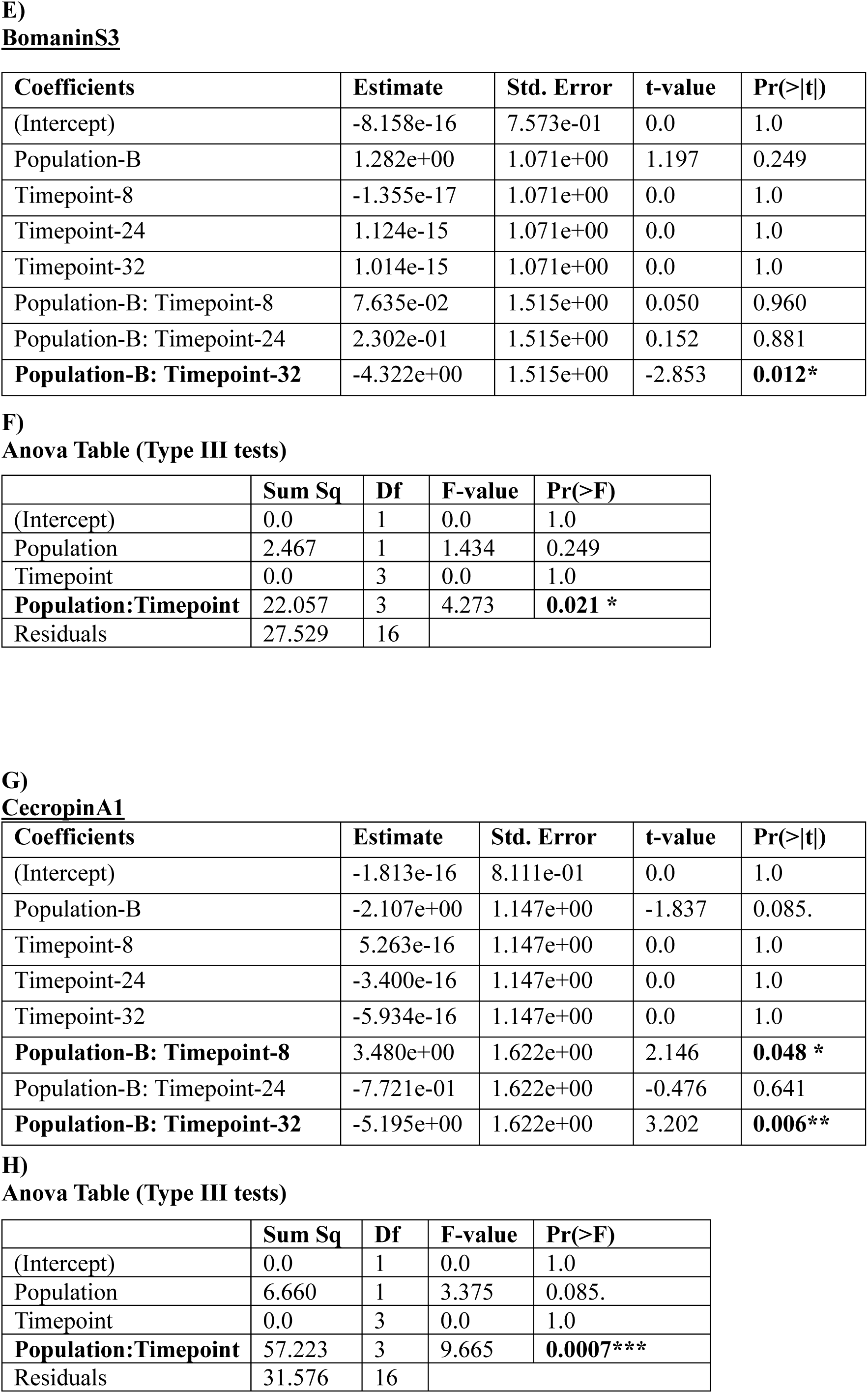
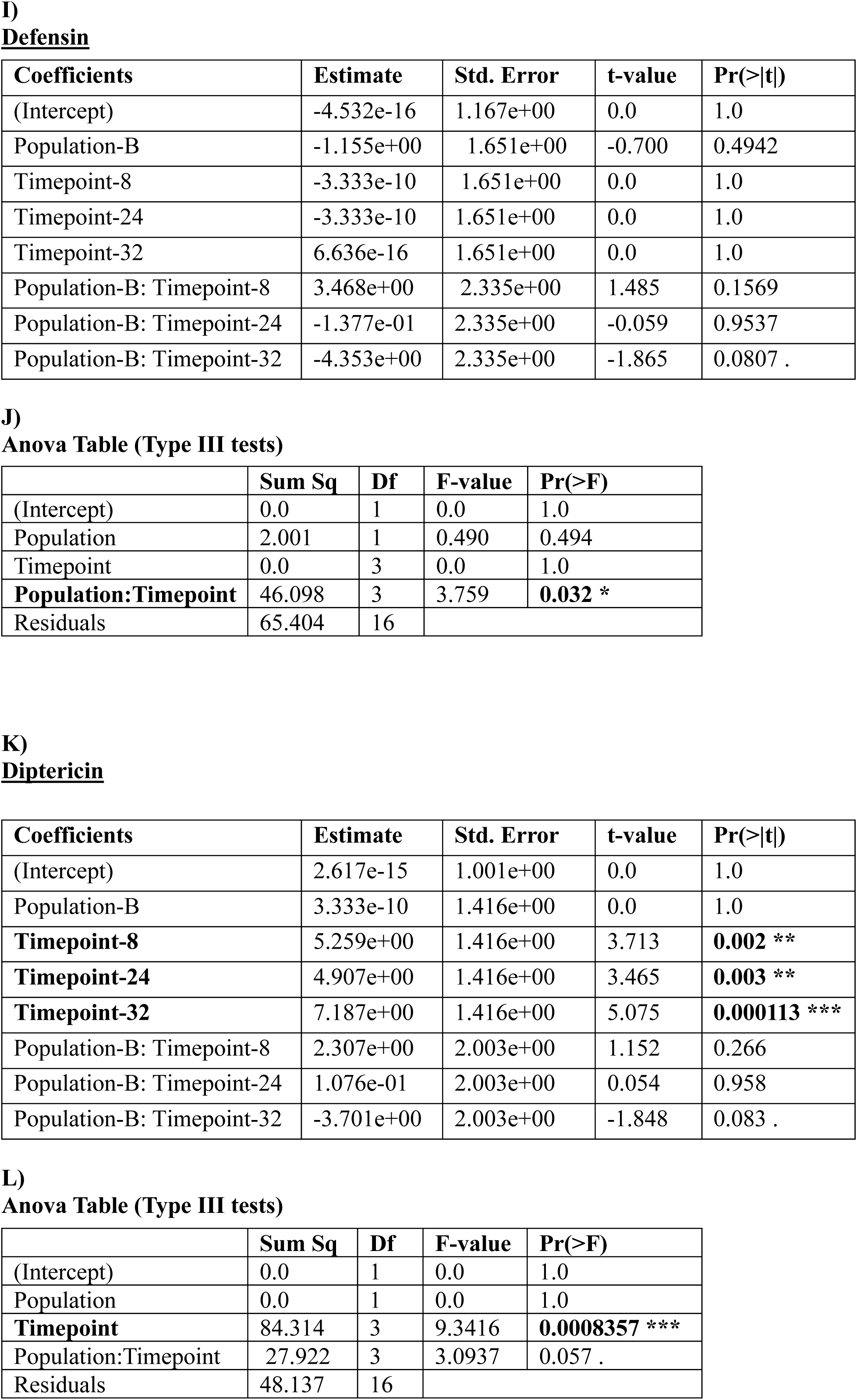
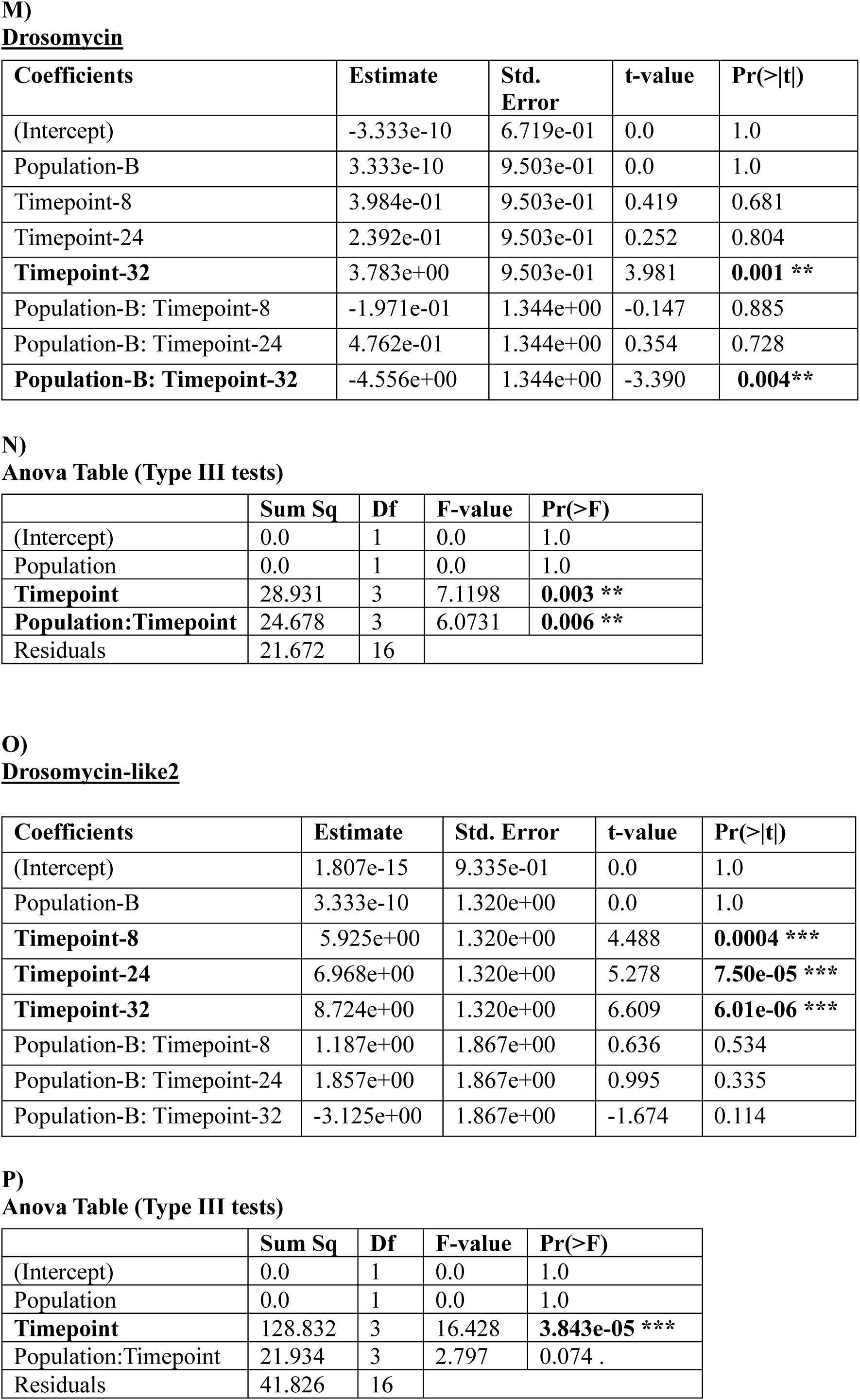
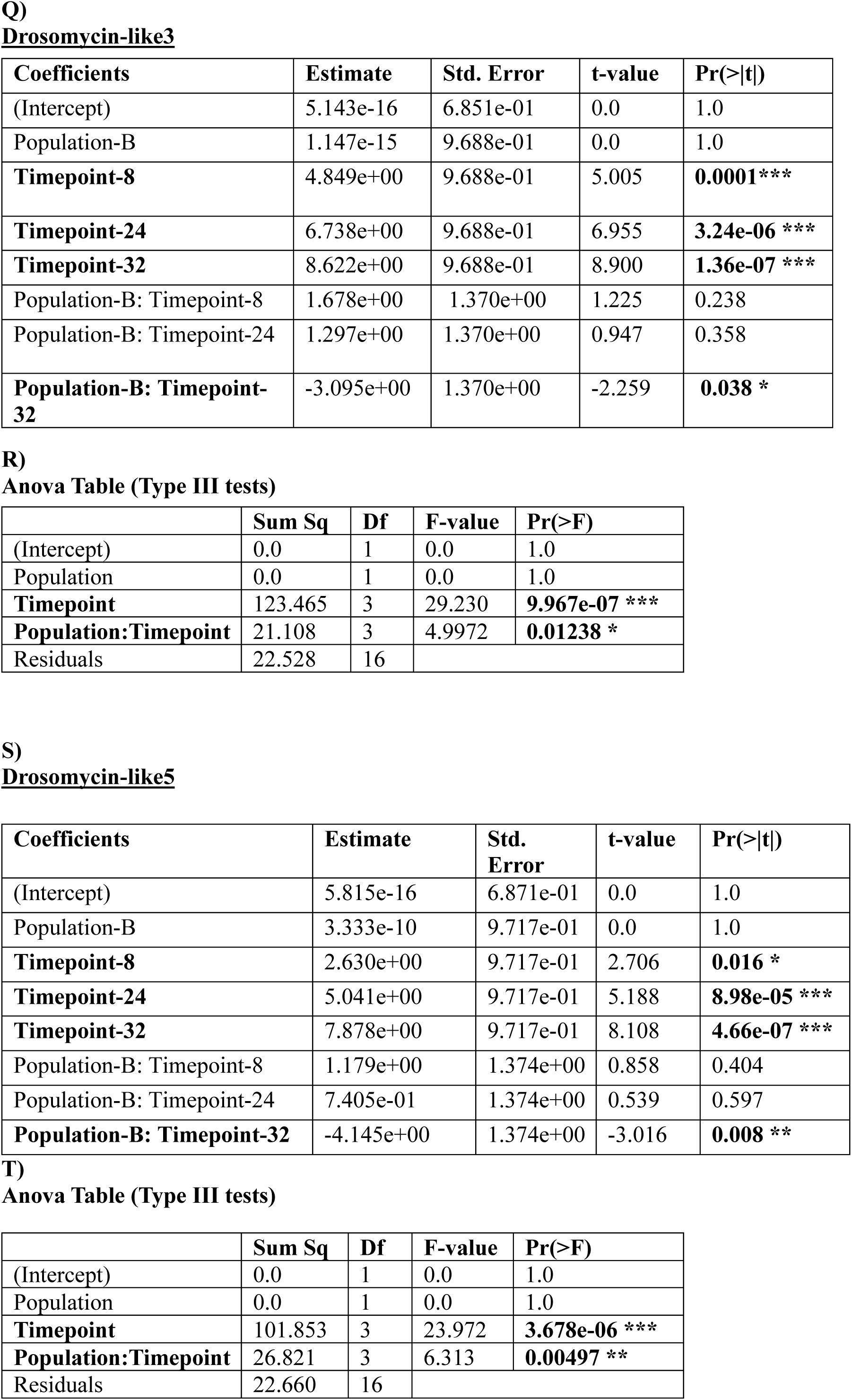
Analysis of relative gene expression levels in guts of females of evolved populations at several timepoints throughout infection (shown in **Fig. 4**). Model estimates and ANOVA estimates for the linear models testing the effects of population and timepoint on expression levels of multiple AMPs during the course of oral bacterial infection.

| Coefficients | Estimate | Std. Error | t-value | Pr(>\|t\|) |
| --- | --- | --- | --- | --- |
| (Intercept) | -3.333e-10 | 8.287e-01 | 0.00 | 1.00 |
| Population-B | 3.333e-10 | 1.172e+00 | 0.00 | 1.00 |
| Timepoint-8 | 5.914e+00 | 1.172e+00 | 5.047 | 0.000119 *** |
| Timepoint-24 | 6.171e+00 | 1.172e+00 | 5.266 | 7.69e-05 *** |
| Timepoint-32 | 7.375e+00 | 1.172e+00 | 6.293 | 1.07e-05 *** |
| Population-B: Timepoint-8 | 3.004e+00 | 1.657e+00 | 1.813 | 0.088712 . |
| Population-B: Timepoint-24 | -1.412e+00 | 1.657e+00 | -0.852 | 0.406965 |
| Population-B: Timepoint-32 | -1.950e+00 | 1.657e+00 | -1.177 | 0.256541 |

B) Anova Table (Type III tests)
|  | Sum Sq | Df | F-value | Pr(>F) |
| --- | --- | --- | --- | --- |
| (Intercept) | 0.0 | 1 | 0.00 | 1.0 |
| Population | 0.0 | 1 | 0.00 | 1.0 |
| **Timepoint** | 98.317 | 3 | 15.908 | **4.647e-05 \*\*\*** |
| **Population:Timepoint** | 22.181 | 3 | 3.589 | **0.03718 \*** |
| Residuals | 32.962 | 16 |  |  |

C) BomaninS2
| Coefficients | Estimate | Std. Error | t-value | Pr(>\|t\|) |
| --- | --- | --- | --- | --- |
| (Intercept) | -5.881e-16 | 8.017e-01 | 0.0 | 1.0 |
| Population-B | 1.358e-15 | 1.134e+00 | 0.0 | 1.0 |
| Timepoint-8 | 1.563e+00 | 1.134e+00 | 1.378 | 0.187 |
| Timepoint-24 | 2.062e+00 | 1.134e+00 | 1.818 | 0.088. |
| **Timepoint-32** | **6.049e+00** | 1.134e+00 | 5.335 | **6.7e-05 \*\*\*** |
| Population-B: Timepoint-8 | 5.023e-01 | 1.603e+00 | 0.313 | 0.758 |
| Population-B: Timepoint-24 | 2.832e-01 | 1.603e+00 | 0.177 | 0.862 |
| **Population-B: Timepoint-32** | -4.928e+00 | 1.603e+00 | -3.074 | **0.007 \*\*** |

D) Anova Table (Type III tests)
|  | Sum Sq | Df | F-value | Pr(>F) |
| --- | --- | --- | --- | --- |
| (Intercept) | 0.0 | 1 | 0.0 | 1.0 |
| Population | 0.0 | 1 | 0.0 | 1.0 |
| **Timepoint** | 59.677 | 3 | 10.3161 | **0.0005071 \*\*\*** |
| **Population:Timepoint** | 30.495 | 3 | 5.2715 | **0.0101493 \*** |
| Residuals | 30.852 | 16 |  |  |

**BomaninS3**
| Coefficients | Estimate | Std. Error | t-value | Pr(>\|t\|) |
| --- | --- | --- | --- | --- |
| (Intercept) | -8.158e-16 | 7.573e-01 | 0.0 | 1.0 |
| Population-B | 1.282e+00 | 1.071e+00 | 1.197 | 0.249 |
| Timepoint-8 | -1.355e-17 | 1.071e+00 | 0.0 | 1.0 |
| Timepoint-24 | 1.124e-15 | 1.071e+00 | 0.0 | 1.0 |
| Timepoint-32 | 1.014e-15 | 1.071e+00 | 0.0 | 1.0 |
| Population-B: Timepoint-8 | 7.635e-02 | 1.515e+00 | 0.050 | 0.960 |
| Population-B: Timepoint-24 | 2.302e-01 | 1.515e+00 | 0.152 | 0.881 |
| **Population-B: Timepoint-32** | -4.322e+00 | 1.515e+00 | -2.853 | **0.012\*** |

**Anova Table (Type III tests)**
|  | Sum Sq | Df | F-value | Pr(>F) |
| --- | --- | --- | --- | --- |
| (Intercept) | 0.0 | 1 | 0.0 | 1.0 |
| Population | 2.467 | 1 | 1.434 | 0.249 |
| Timepoint | 0.0 | 3 | 0.0 | 1.0 |
| **Population:Timepoint** | 22.057 | 3 | 4.273 | **0.021 \*** |
| Residuals | 27.529 | 16 |  |  |

**CecropinA1**
| Coefficients | Estimate | Std. Error | t-value | Pr(>\|t\|) |
| --- | --- | --- | --- | --- |
| (Intercept) | -1.813e-16 | 8.111e-01 | 0.0 | 1.0 |
| Population-B | -2.107e+00 | 1.147e+00 | -1.837 | 0.085. |
| Timepoint-8 | 5.263e-16 | 1.147e+00 | 0.0 | 1.0 |
| Timepoint-24 | -3.400e-16 | 1.147e+00 | 0.0 | 1.0 |
| Timepoint-32 | -5.934e-16 | 1.147e+00 | 0.0 | 1.0 |
| **Population-B: Timepoint-8** | 3.480e+00 | 1.622e+00 | 2.146 | **0.048 \*** |
| Population-B: Timepoint-24 | -7.721e-01 | 1.622e+00 | -0.476 | 0.641 |
| **Population-B: Timepoint-32** | -5.195e+00 | 1.622e+00 | 3.202 | **0.006\*\*** |

**Anova Table (Type III tests)**
|  | Sum Sq | Df | F-value | Pr(>F) |
| --- | --- | --- | --- | --- |
| (Intercept) | 0.0 | 1 | 0.0 | 1.0 |
| Population | 6.660 | 1 | 3.375 | 0.085. |
| Timepoint | 0.0 | 3 | 0.0 | 1.0 |
| **Population:Timepoint** | 57.223 | 3 | 9.665 | **0.0007\*\*\*** |
| Residuals | 31.576 | 16 |  |  |

**Defensin**
| Coefficients | Estimate | Std. Error | t-value | Pr(>\|t\|) |
| --- | --- | --- | --- | --- |
| (Intercept) | -4.532e-16 | 1.167e+00 | 0.0 | 1.0 |
| Population-B | -1.155e+00 | 1.651e+00 | -0.700 | 0.4942 |
| Timepoint-8 | -3.333e-10 | 1.651e+00 | 0.0 | 1.0 |
| Timepoint-24 | -3.333e-10 | 1.651e+00 | 0.0 | 1.0 |
| Timepoint-32 | 6.636e-16 | 1.651e+00 | 0.0 | 1.0 |
| Population-B: Timepoint-8 | 3.468e+00 | 2.335e+00 | 1.485 | 0.1569 |
| Population-B: Timepoint-24 | -1.377e-01 | 2.335e+00 | -0.059 | 0.9537 |
| Population-B: Timepoint-32 | -4.353e+00 | 2.335e+00 | -1.865 | 0.0807 . |

**Anova Table (Type III tests)**
|  | Sum Sq | Df | F-value | Pr(>F) |
| --- | --- | --- | --- | --- |
| (Intercept) | 0.0 | 1 | 0.0 | 1.0 |
| Population | 2.001 | 1 | 0.490 | 0.494 |
| Timepoint | 0.0 | 3 | 0.0 | 1.0 |
| **Population:Timepoint** | 46.098 | 3 | 3.759 | **0.032 \*** |
| Residuals | 65.404 | 16 |  |  |

**Diptericin**
| Coefficients | Estimate | Std. Error | t-value | Pr(>\|t\|) |
| --- | --- | --- | --- | --- |
| (Intercept) | 2.617e-15 | 1.001e+00 | 0.0 | 1.0 |
| Population-B | 3.333e-10 | 1.416e+00 | 0.0 | 1.0 |
| **Timepoint-8** | 5.259e+00 | 1.416e+00 | 3.713 | **0.002 \*\*** |
| **Timepoint-24** | 4.907e+00 | 1.416e+00 | 3.465 | **0.003 \*\*** |
| **Timepoint-32** | 7.187e+00 | 1.416e+00 | 5.075 | **0.000113 \*\*\*** |
| Population-B: Timepoint-8 | 2.307e+00 | 2.003e+00 | 1.152 | 0.266 |
| Population-B: Timepoint-24 | 1.076e-01 | 2.003e+00 | 0.054 | 0.958 |
| Population-B: Timepoint-32 | -3.701e+00 | 2.003e+00 | -1.848 | 0.083 . |

**Anova Table (Type III tests)**
|  | Sum Sq | Df | F-value | Pr(>F) |
| --- | --- | --- | --- | --- |
| (Intercept) | 0.0 | 1 | 0.0 | 1.0 |
| Population | 0.0 | 1 | 0.0 | 1.0 |
| **Timepoint** | 84.314 | 3 | 9.3416 | **0.0008357 \*\*\*** |
| Population:Timepoint | 27.922 | 3 | 3.0937 | 0.057 . |
| Residuals | 48.137 | 16 |  |  |

**Drosomycin**
| Coefficients | Estimate | Std. Error | t-value | Pr(>\|t\|) |
| --- | --- | --- | --- | --- |
| (Intercept) | -3.333e-10 | 6.719e-01 | 0.0 | 1.0 |
| Population-B | 3.333e-10 | 9.503e-01 | 0.0 | 1.0 |
| Timepoint-8 | 3.984e-01 | 9.503e-01 | 0.419 | 0.681 |
| Timepoint-24 | 2.392e-01 | 9.503e-01 | 0.252 | 0.804 |
| **Timepoint-32** | 3.783e+00 | 9.503e-01 | 3.981 | **0.001 \*\*** |
| Population-B: Timepoint-8 | -1.971e-01 | 1.344e+00 | -0.147 | 0.885 |
| Population-B: Timepoint-24 | 4.762e-01 | 1.344e+00 | 0.354 | 0.728 |
| **Population-B: Timepoint-32** | -4.556e+00 | 1.344e+00 | -3.390 | **0.004\*\*** |

**Anova Table (Type III tests)**
|  | Sum Sq | Df | F-value | Pr(>F) |
| --- | --- | --- | --- | --- |
| (Intercept) | 0.0 | 1 | 0.0 | 1.0 |
| Population | 0.0 | 1 | 0.0 | 1.0 |
| **Timepoint** | 28.931 | 3 | 7.1198 | **0.003 \*\*** |
| **Population:Timepoint** | 24.678 | 3 | 6.0731 | **0.006 \*\*** |
| Residuals | 21.672 | 16 |  |  |

**Drosomycin-like2**
| Coefficients | Estimate | Std. Error | t-value | Pr(>\|t\|) |
| --- | --- | --- | --- | --- |
| (Intercept) | 1.807e-15 | 9.335e-01 | 0.0 | 1.0 |
| Population-B | 3.333e-10 | 1.320e+00 | 0.0 | 1.0 |
| **Timepoint-8** | 5.925e+00 | 1.320e+00 | 4.488 | **0.0004 \*\*\*** |
| **Timepoint-24** | 6.968e+00 | 1.320e+00 | 5.278 | **7.50e-05 \*\*\*** |
| **Timepoint-32** | 8.724e+00 | 1.320e+00 | 6.609 | **6.01e-06 \*\*\*** |
| Population-B: Timepoint-8 | 1.187e+00 | 1.867e+00 | 0.636 | 0.534 |
| Population-B: Timepoint-24 | 1.857e+00 | 1.867e+00 | 0.995 | 0.335 |
| Population-B: Timepoint-32 | -3.125e+00 | 1.867e+00 | -1.674 | 0.114 |

**Anova Table (Type III tests)**
|  | Sum Sq | Df | F-value | Pr(>F) |
| --- | --- | --- | --- | --- |
| (Intercept) | 0.0 | 1 | 0.0 | 1.0 |
| Population | 0.0 | 1 | 0.0 | 1.0 |
| **Timepoint** | 128.832 | 3 | 16.428 | **3.843e-05 \*\*\*** |
| Population:Timepoint | 21.934 | 3 | 2.797 | 0.074 . |
| Residuals | 41.826 | 16 |  |  |

**Drosomycin-like3**
| Coefficients | Estimate | Std. Error | t-value | Pr(>\|t\|) |
| --- | --- | --- | --- | --- |
| (Intercept) | 5.143e-16 | 6.851e-01 | 0.0 | 1.0 |
| Population-B | 1.147e-15 | 9.688e-01 | 0.0 | 1.0 |
| **Timepoint-8** | 4.849e+00 | 9.688e-01 | 5.005 | **0.0001\*\*\*** |
| **Timepoint-24** | 6.738e+00 | 9.688e-01 | 6.955 | **3.24e-06 \*\*\*** |
| **Timepoint-32** | 8.622e+00 | 9.688e-01 | 8.900 | **1.36e-07 \*\*\*** |
| Population-B: Timepoint-8 | 1.678e+00 | 1.370e+00 | 1.225 | 0.238 |
| Population-B: Timepoint-24 | 1.297e+00 | 1.370e+00 | 0.947 | 0.358 |
| **Population-B: Timepoint-32** | -3.095e+00 | 1.370e+00 | -2.259 | **0.038 \*** |

**Anova Table (Type III tests)**
|  | Sum Sq | Df | F-value | Pr(>F) |
| --- | --- | --- | --- | --- |
| (Intercept) | 0.0 | 1 | 0.0 | 1.0 |
| Population | 0.0 | 1 | 0.0 | 1.0 |
| **Timepoint** | 123.465 | 3 | 29.230 | **9.967e-07 \*\*\*** |
| **Population:Timepoint** | 21.108 | 3 | 4.9972 | **0.01238 \*** |
| Residuals | 22.528 | 16 |  |  |

**Drosomycin-like5**
| Coefficients | Estimate | Std. Error | t-value | Pr(>\|t\|) |
| --- | --- | --- | --- | --- |
| (Intercept) | 5.815e-16 | 6.871e-01 | 0.0 | 1.0 |
| Population-B | 3.333e-10 | 9.717e-01 | 0.0 | 1.0 |
| **Timepoint-8** | 2.630e+00 | 9.717e-01 | 2.706 | **0.016 \*** |
| **Timepoint-24** | 5.041e+00 | 9.717e-01 | 5.188 | **8.98e-05 \*\*\*** |
| **Timepoint-32** | 7.878e+00 | 9.717e-01 | 8.108 | **4.66e-07 \*\*\*** |
| Population-B: Timepoint-8 | 1.179e+00 | 1.374e+00 | 0.858 | 0.404 |
| Population-B: Timepoint-24 | 7.405e-01 | 1.374e+00 | 0.539 | 0.597 |
| **Population-B: Timepoint-32** | -4.145e+00 | 1.374e+00 | -3.016 | **0.008 \*\*** |

**Supplementary Table 6. Analysis of relative gene expression levels in guts of females of evolved populations at several timepoints throughout infection (shown in Fig. 4).**
|  | Sum Sq | Df | F-value | Pr(>F) |
| --- | --- | --- | --- | --- |
| (Intercept) | 0.0 | 1 | 0.0 | 1.0 |
| Population | 0.0 | 1 | 0.0 | 1.0 |
| **Timepoint** | 101.853 | 3 | 23.972 | **3.678e-06 \*\*\*** |
| **Population:Timepoint** | 26.821 | 3 | 6.313 | **0.00497 \*\*** |
| Residuals | 22.660 | 16 |  |  |

**Supplementary Table 7.**
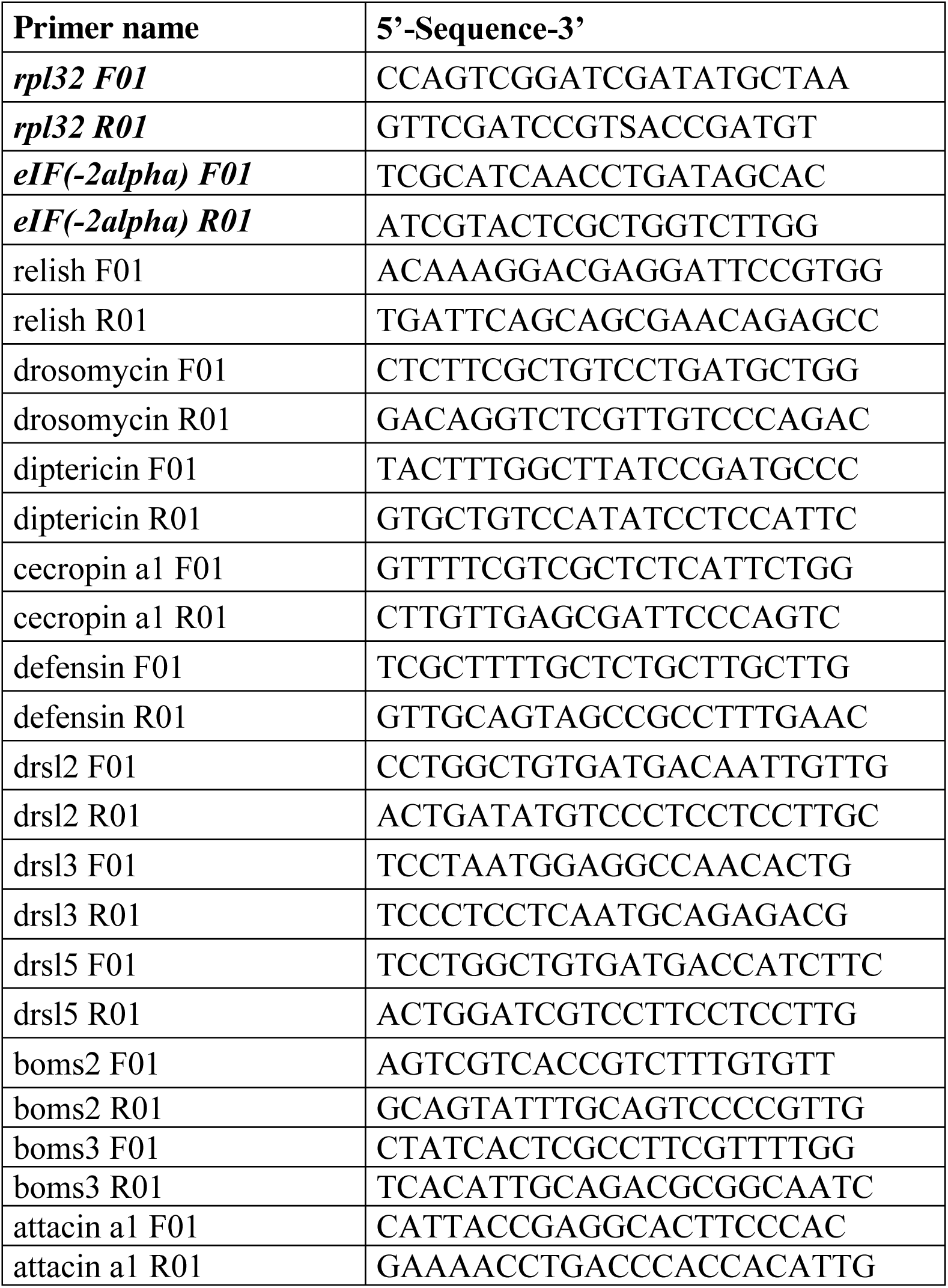
Primer sequences used in Real-Time quantitative PCRs (shown in Fig. 4). The first four genes listed (shown in bold and italics) were used as reference (house-keeping) for relative quantifications.

**Supplementary Table 8.** Statistical analysis of survival of evolved populations after priming with heat-killed bacteria (shown in Fig. 5). A) Model estimates and B) ANOVA estimates for survival analysis testing the effect of priming on infection in Control and BactOral.

| **Fixed coefficients** | **coef** | **exp(coef)** | **se(coef)** | **z** | **p** |
| --- | --- | --- | --- | --- | --- |
| **Population-B** | -1.065 | 0.345 | 0.156 | -6.82 | **8.8e-12** |
| **Treatment-HK** | -1.324 | 0.266 | 0.132 | -10.01 | **0.0e+00** |
| Sex-m | -0.195 | 0.823 | 0.130 | -1.50 | 1.3e-01 |
| Population-B:Treatment-HK | -0.115 | 0.891 | 0.239 | -0.48 | 6.3e-01 |
| **Population-B: Sex-m** | -0.792 | 0.453 | 0.256 | -3.09 | **2.0e-03** |
| **Treatment-HK: Sex-m** | 0.382 | 1.465 | 0.184 | 2.08 | **3.8e-02** |
| Population-B:Treatment-HK:Sex-m | -0.172 | 0.842 | 0.391 | -0.44 | 6.6e-01 |
| **Random effects** |  |  |  |  |  |
| **Group** | **Variable** | **Std Dev** | **Variance** |  |  |
| Block/Replicate | (Intercept) | 0.152 | 0.023 |  |  |
| Block | (Intercept) | 0.173 | 0.030 |  |  |

**Supplementary Table 8. Statistical analysis of survival of evolved populations after priming with heat-killed bacteria (shown in Fig. 5).**
|  | **Df** | **Chisq** | **Pr(>Chisq)** |
| --- | --- | --- | --- |
| **Population** | 1 | 46.577 | **8.808e-12 \*\*\*** |
| **Treatment** | 1 | 100.214 | **< 2.2e-16 \*\*\*** |
| Sex | 1 | 2.238 | 0.135 |
| Population:Treatment | 1 | 0.233 | 0.639 |
| **Population:Sex** | 1 | 9.553 | **0.002 \*\*** |
| **Treatment:Sex** | 1 | 4.308 | **0.038 \*** |
| Population:Treatment:Sex | 1 | 0.194 | 0.659 |

**Supplementary Table 9.**
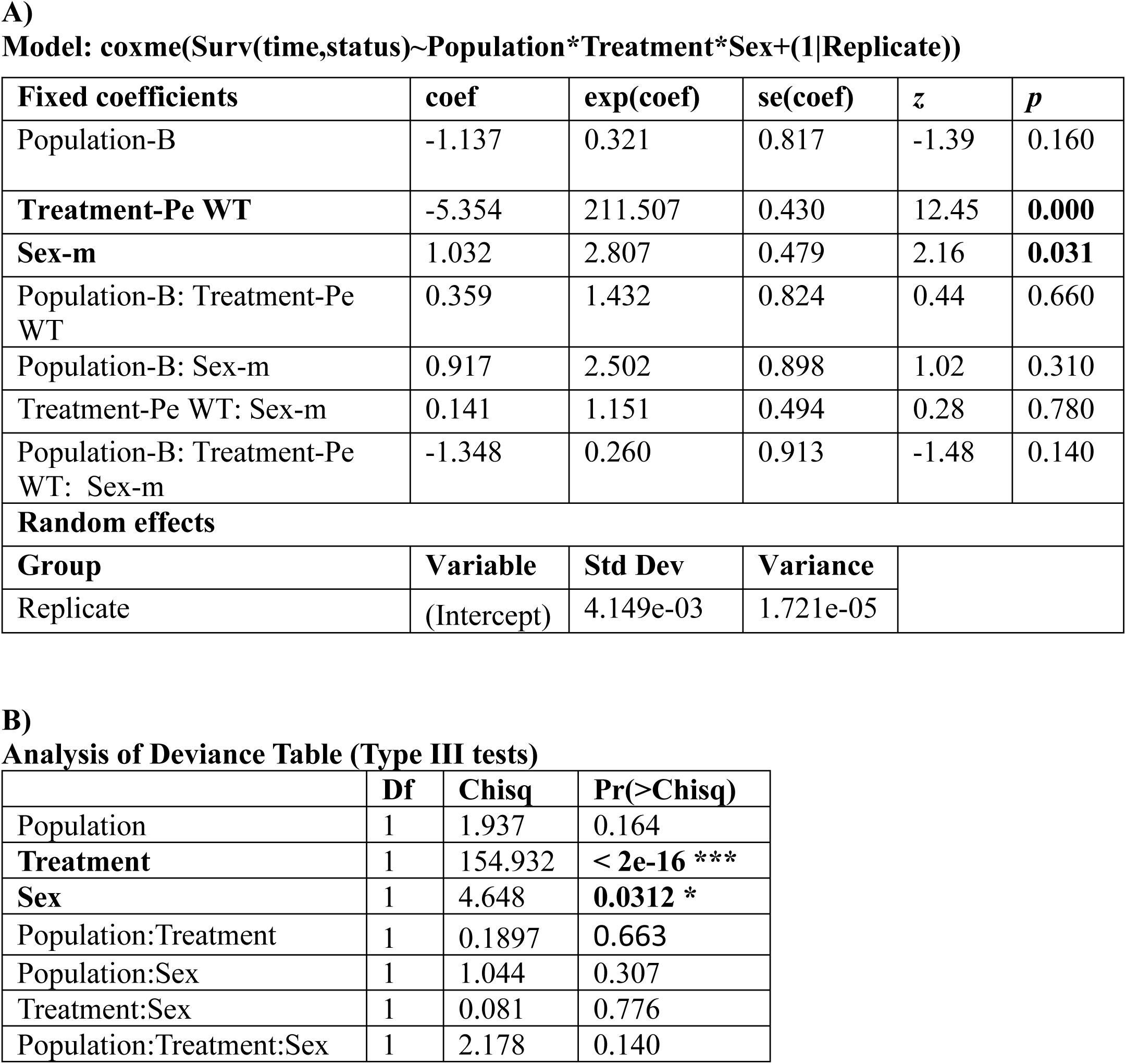
Statistical analysis of survival of evolved populations after prolonged exposure to heat-killed bacteria (shown in Fig. 6). A) Model estimates and B) ANOVA estimates for survival analysis testing the effects of population, treatment and sex on survival to prolonged exposure to heat-killed *P. entomophila*, for 14 days.

| Fixed coefficients | coef | exp(coef) | se(coef) | z | p |
| --- | --- | --- | --- | --- | --- |
| Population-B | -1.137 | 0.321 | 0.817 | -1.39 | 0.160 |
| **Treatment-Pe WT** | -5.354 | 211.507 | 0.430 | 12.45 | **0.000** |
| **Sex-m** | 1.032 | 2.807 | 0.479 | 2.16 | **0.031** |
| Population-B: Treatment-Pe WT | 0.359 | 1.432 | 0.824 | 0.44 | 0.660 |
| Population-B: Sex-m | 0.917 | 2.502 | 0.898 | 1.02 | 0.310 |
| Treatment-Pe WT: Sex-m | 0.141 | 1.151 | 0.494 | 0.28 | 0.780 |
| Population-B: Treatment-Pe WT: Sex-m | -1.348 | 0.260 | 0.913 | -1.48 | 0.140 |
| **Random effects** |  |  |  |  |  |
| Group | Variable | Std Dev | Variance |  |  |
| Replicate | (Intercept) | 4.149e-03 | 1.721e-05 |  |  |

**Supplementary Table 9. Statistical analysis of survival of evolved populations after prolonged exposure to heat-killed bacteria (shown in Fig. 6).**
|  | Df | Chisq | Pr(>Chisq) |
| --- | --- | --- | --- |
| Population | 1 | 1.937 | 0.164 |
| **Treatment** | 1 | 154.932 | **< 2e-16 \*\*\*** |
| **Sex** | 1 | 4.648 | **0.0312 \*** |
| Population:Treatment | 1 | 0.1897 | 0.663 |
| Population:Sex | 1 | 1.044 | 0.307 |
| Treatment:Sex | 1 | 0.081 | 0.776 |
| Population:Treatment:Sex | 1 | 2.178 | 0.140 |

